## Supplementary Material for "Selecting causal risk factors from high-throughput experiments using multivariable Mendelian randomization"

### Contents

**1 Methods** **3**
1.1 Supplementary directed acyclic graph . . . . . . . . . . . . . . . 3

**2 Simulation** **4**
2.1 Supplementary Figures: Simulation results on NMR metabolite data . . . . . . . . . . . . . . . . . . . . . . . . . . . . . . . . . . 4
2.2 Supplementary Figures: Simulation results on blood cell trait data 11

**3 Application: Metabolites as risk factors for age-related macular degeneration** **15**
3.1 Supplementary Tables . . . . . . . . . . . . . . . . . . . . . . . . 15
3.2 Supplementary Figures . . . . . . . . . . . . . . . . . . . . . . . . 27

14 Causal effect estimate for setting A for the blood cell traits ($d = 33$) 13
15 Causal effect estimate for setting B for the blood cell traits ($d = 33$) 14

16 Diagnostic plots of the predicted associations with AMD against the observed associations with AMD . . . . . . . . . . . . . . . . 27
17 Scatterplot of association with A) XL.HDL.C B) L.HDL.C on the $x$-axis against the association with AMD $y$-axis after excluding the *LIPC*, *FUT2* and *APOE* gene regions. The model-averaged causal effect (MACE) of each risk factor on AMD is marked in red. . . . . . . . . . . . . . . . . . . . . . . . . . . . . . . . . . 28
18 Diagnostic plots of the predicted associations with AMD against the observed associations with AMD after excluding the *LIPC, FUT2* and *APOE* gene regions . . . . . . . . . . . . . . . . . . . 29
19 Diagnostic plots of the predicted associations with AMD against the observed associations with AMD after excluding the *LIPC, FUT2* and *APOE* gene regions . . . . . . . . . . . . . . . . . . . 30

### 1 Methods

#### 1.1 Supplementary directed acyclic graph

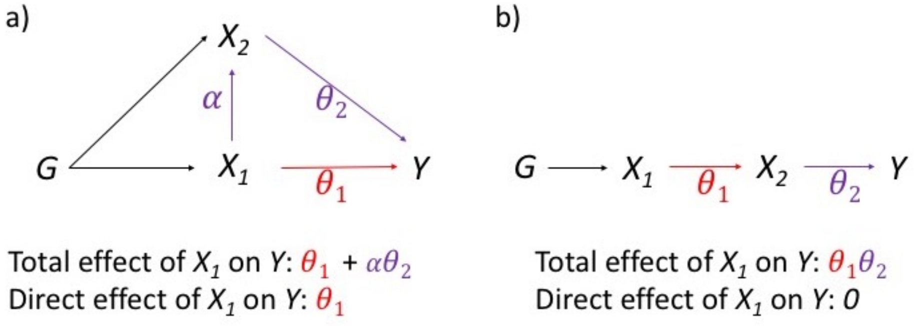

Figure 1: Directed acyclic graph to illustrate the difference between total and direct effect in two scenarios: a) mediation effect, where the risk factor $X_1$ has a direct and an indirect effect via the mediator $X_2$ on the outcome $Y$ and b) signalling cascade where the effect of $X_1$ on the outcome is entirely mediated by $X_2$.

### 2 Simulation

#### 2.1 Supplementary Figures: Simulation results on NMR metabolite data

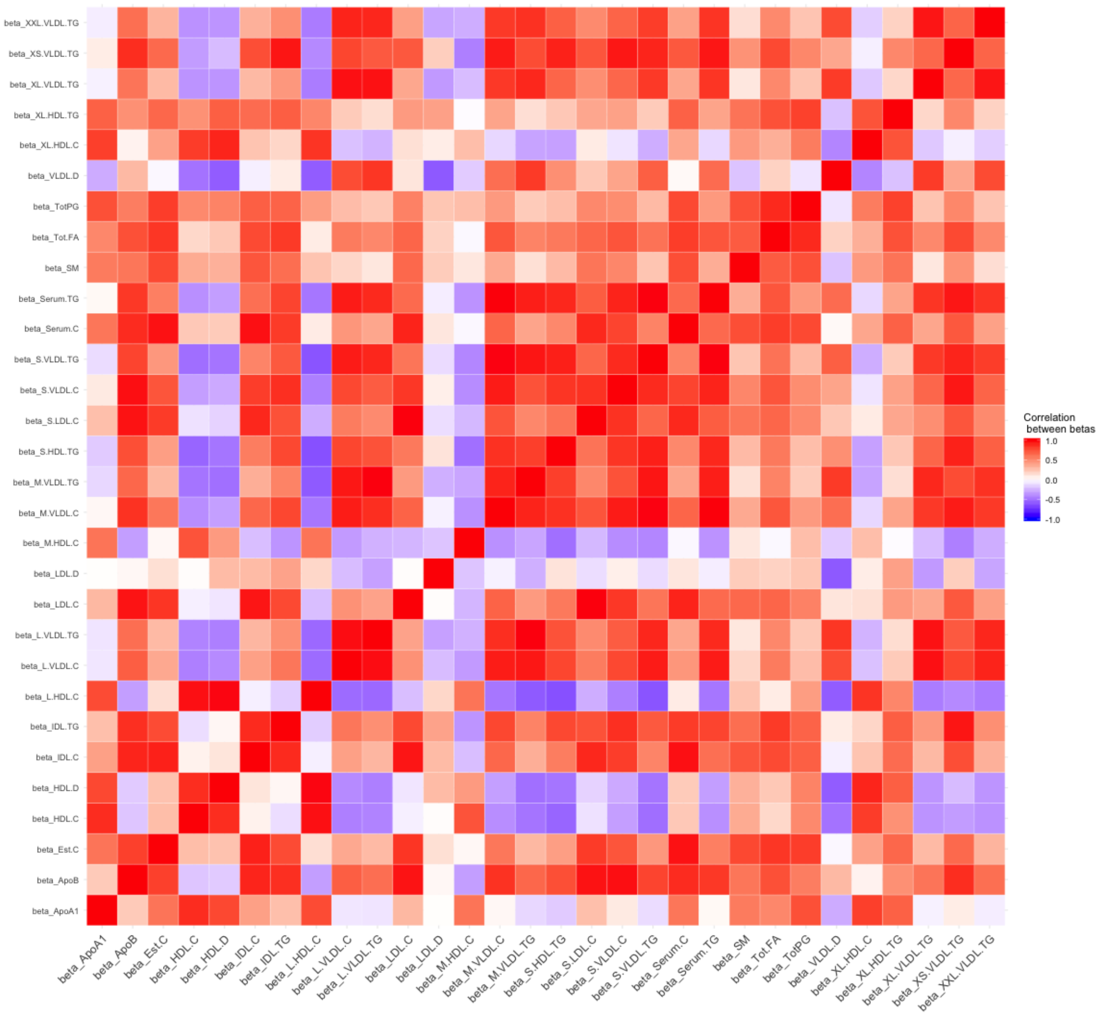

Figure 2: Genetic correlation between metabolite measurements based on the $n = 148$ genetic variants used as instrumental variables.

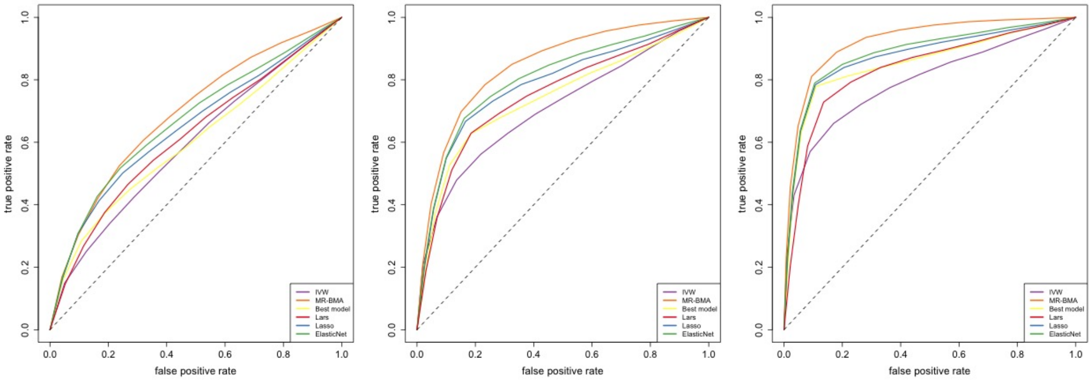

Figure 3: Receiver operating characteristic (ROC) curve for setting A including a small number of risk factors ($d = 12$) of which four are true positive effects. Proportion of variance explained is set to 0.1 (left) 0.3 (middle) and 0.5 (right).

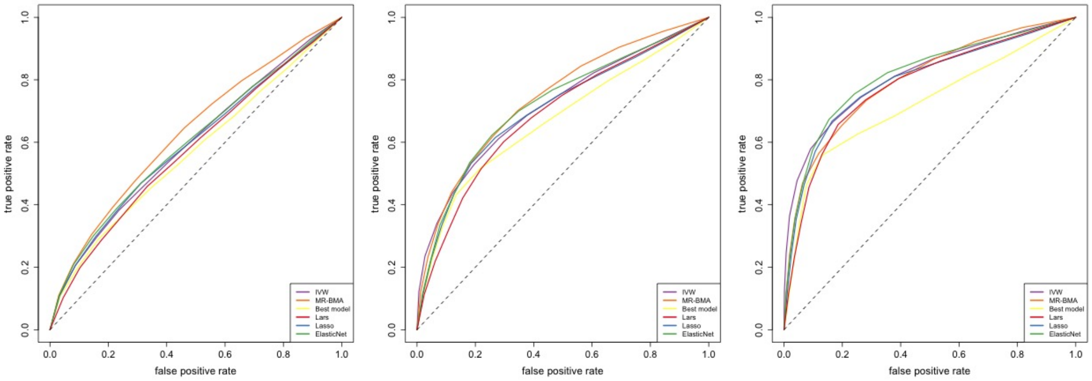

Figure 4: Receiver operating characteristic (ROC) curve for setting B including a small number of risk factors ($d = 12$) of which eight are true positive effects (four postive and four negative effect direction). Proportion of variance explained is set to 0.1 (left) 0.3 (middle) and 0.5 (right).

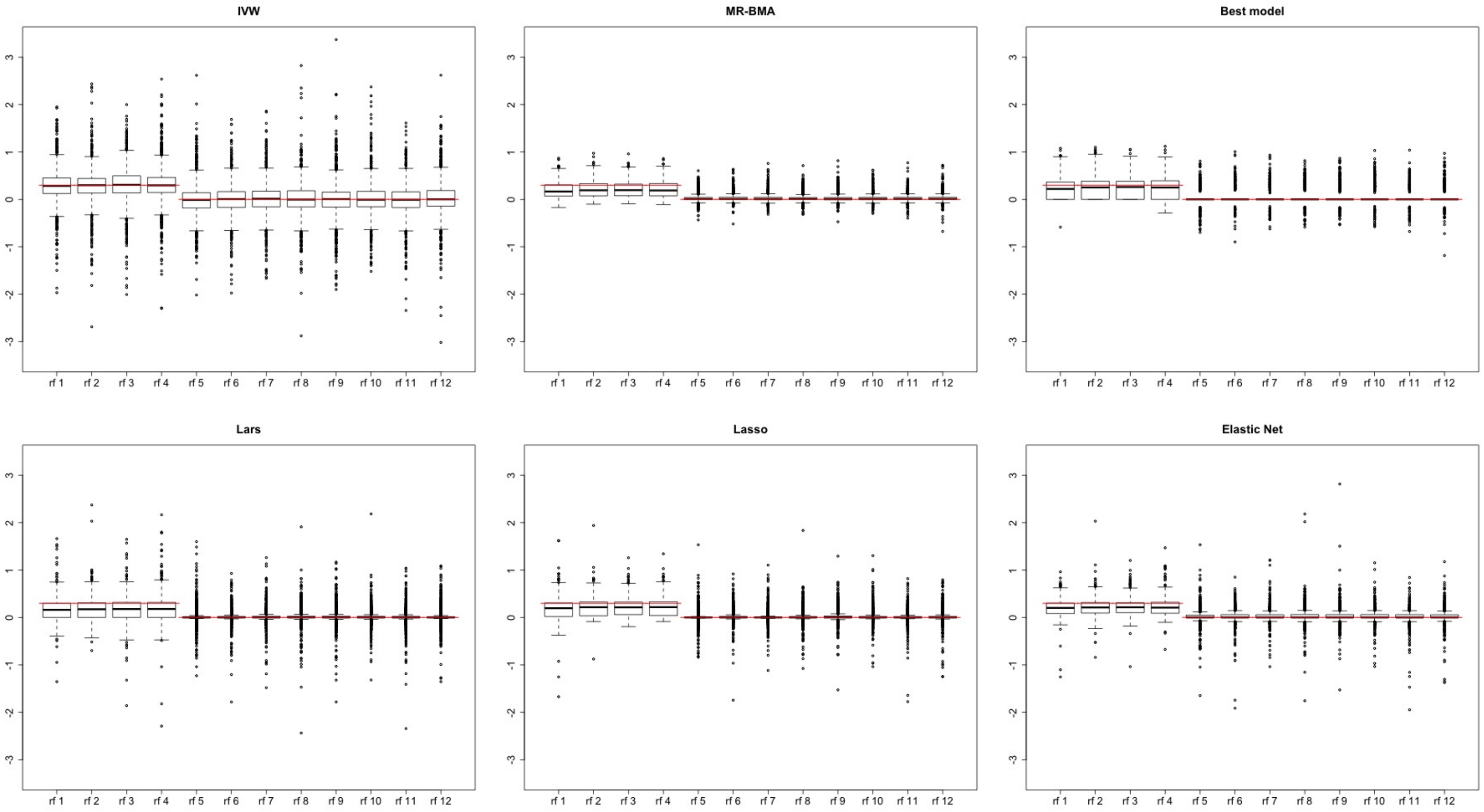

Figure 5: Boxplots of the causal effect estimates for setting A including a small number of risk factors ($d = 12$) of which the first four are true positive effects. The true causal effects are marked in red. From top left to bottom right are the competing approaches: IVW, MR-BMA, best model, Lars, Lasso, and Elastic Net. Proportion of variance explained is set to 0.3.

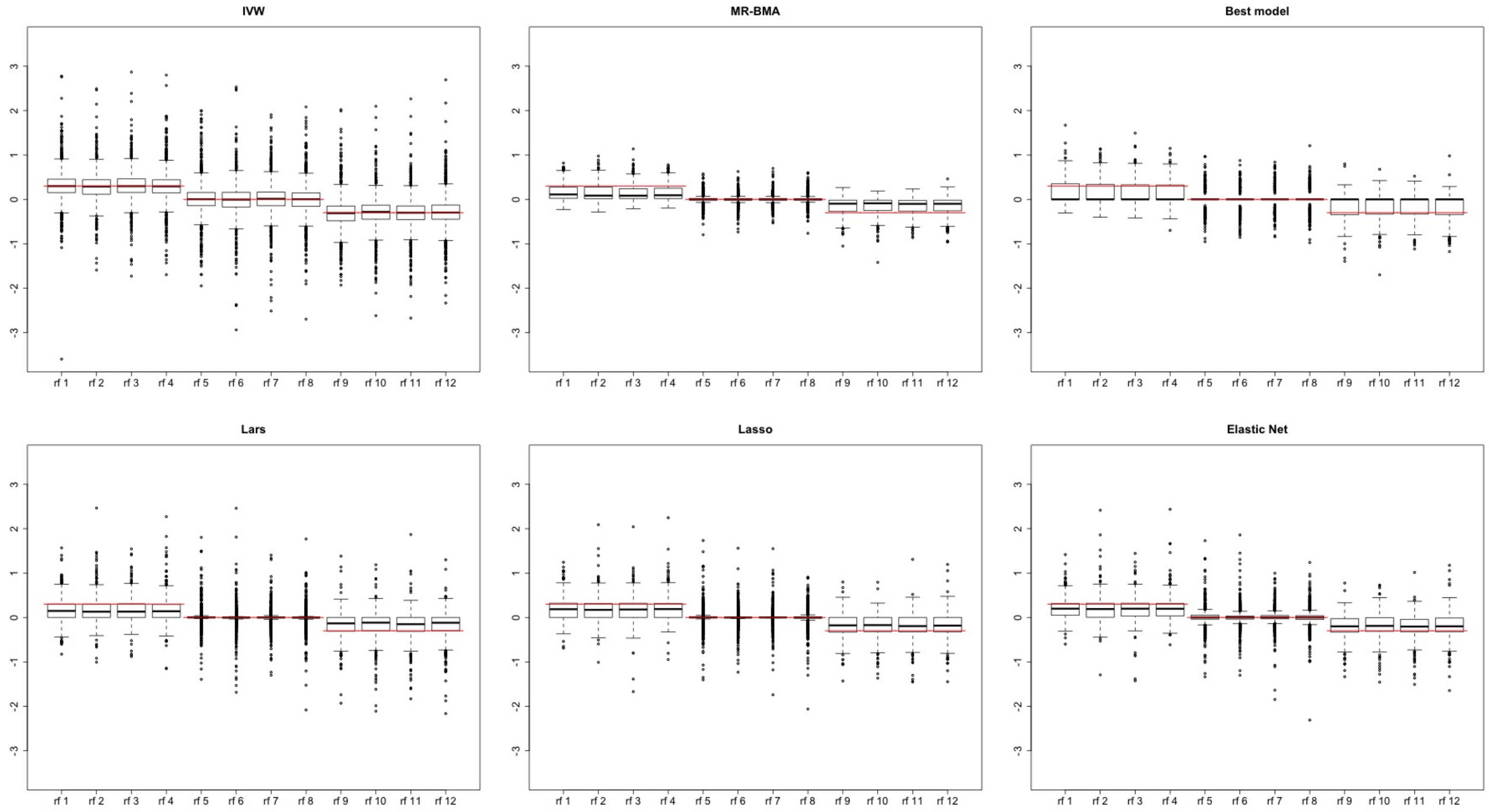

Figure 6: Boxplots of the causal effect estimates for setting B including a small number of risk factors ($d = 12$) of which the first four have a positive and final four have a negative causal effect. The true causal effects are marked in red. From top left to bottom right are the competing approaches: IVW, MR-BMA, best model, Lars, Lasso, and Elastic Net. Proportion of variance explained is set to 0.3.

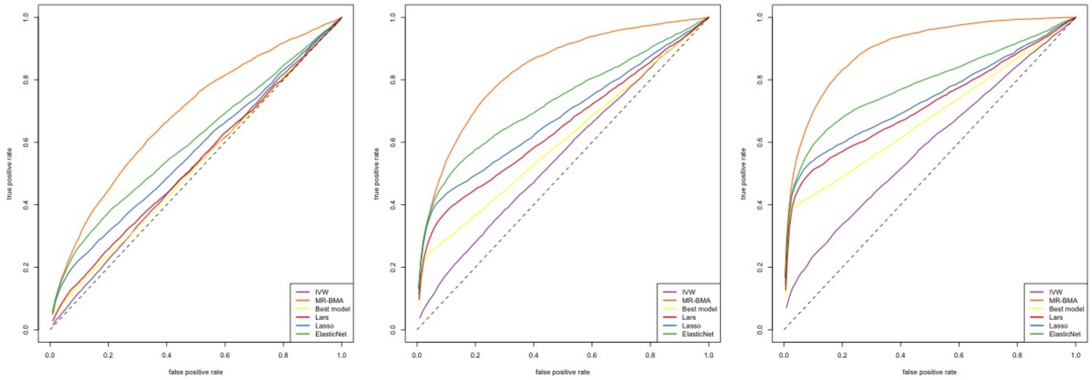

Figure 7: Receiver operating characteristic (ROC) curve for setting A including a large number of risk factors ($d = 92$) of which four are true positive effects. Proportion of variance explained is set to 0.1 (left) 0.3 (middle) and 0.5 (right).

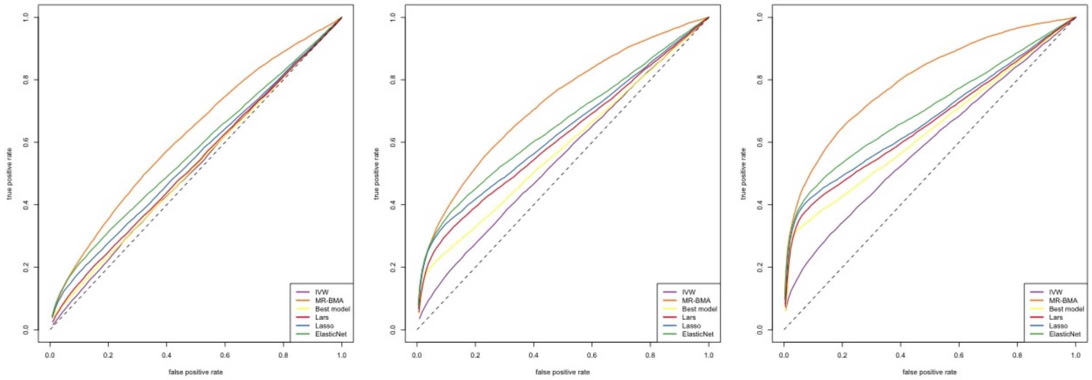

Figure 8: Receiver operating characteristic (ROC) curve for setting B including a large number of risk factors ($d = 92$) of which eight are true positive effects (four postive and four negative effect direction). Proportion of variance explained is set to 0.1 (left) 0.3 (middle) and 0.5 (right).

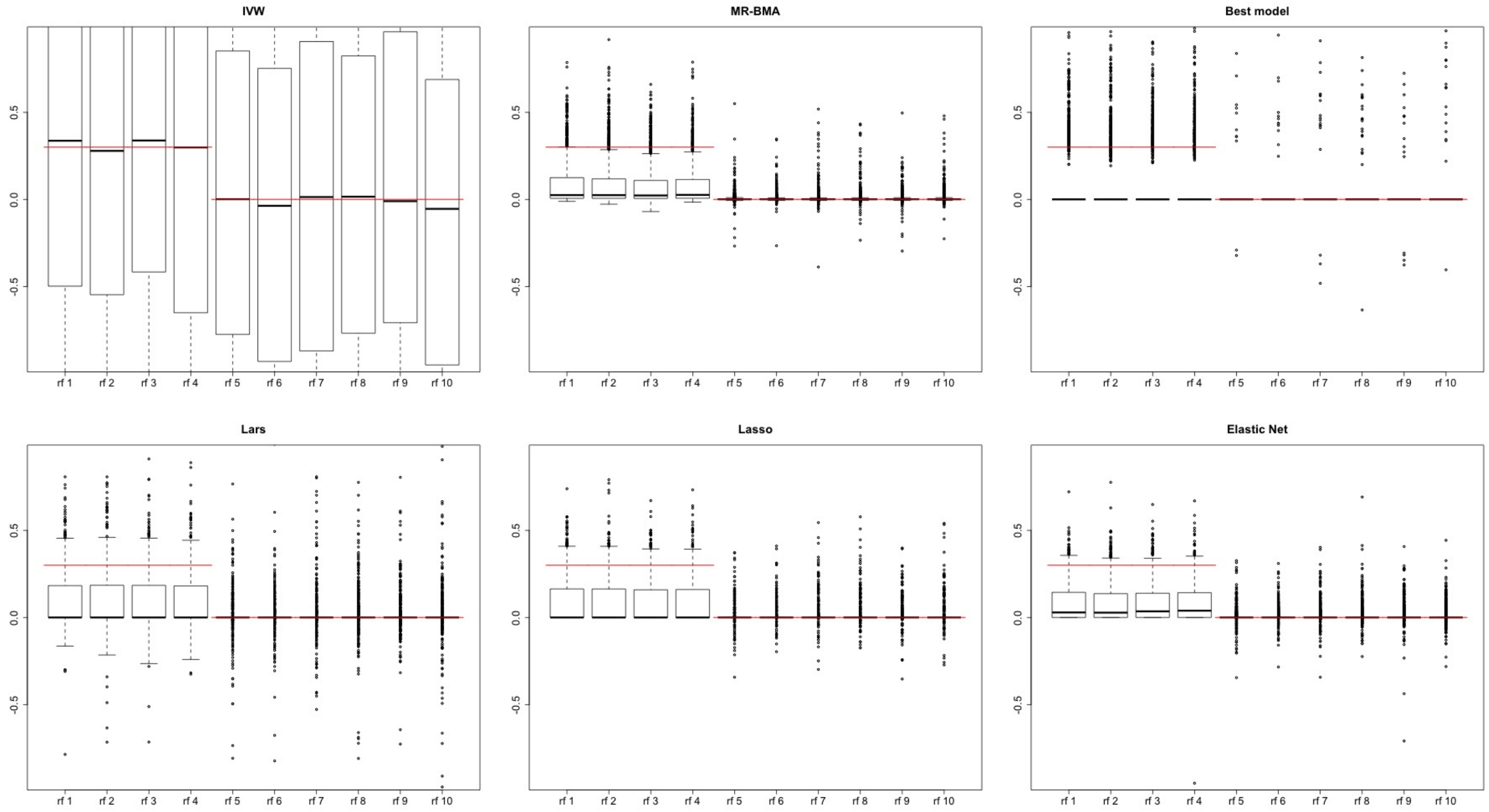

Figure 9: Boxplots of the causal effect estimates for setting A including a large number of risk factors ($d = 92$) of which the first four are true positive effects. Risk factors 11 to 92 are omitted. The true causal effects are marked in red. From top left to bottom right are the competing approaches: IVW, MR-BMA, best model, Lars, Lasso, and Elastic Net. Proportion of variance explained is set to 0.3.

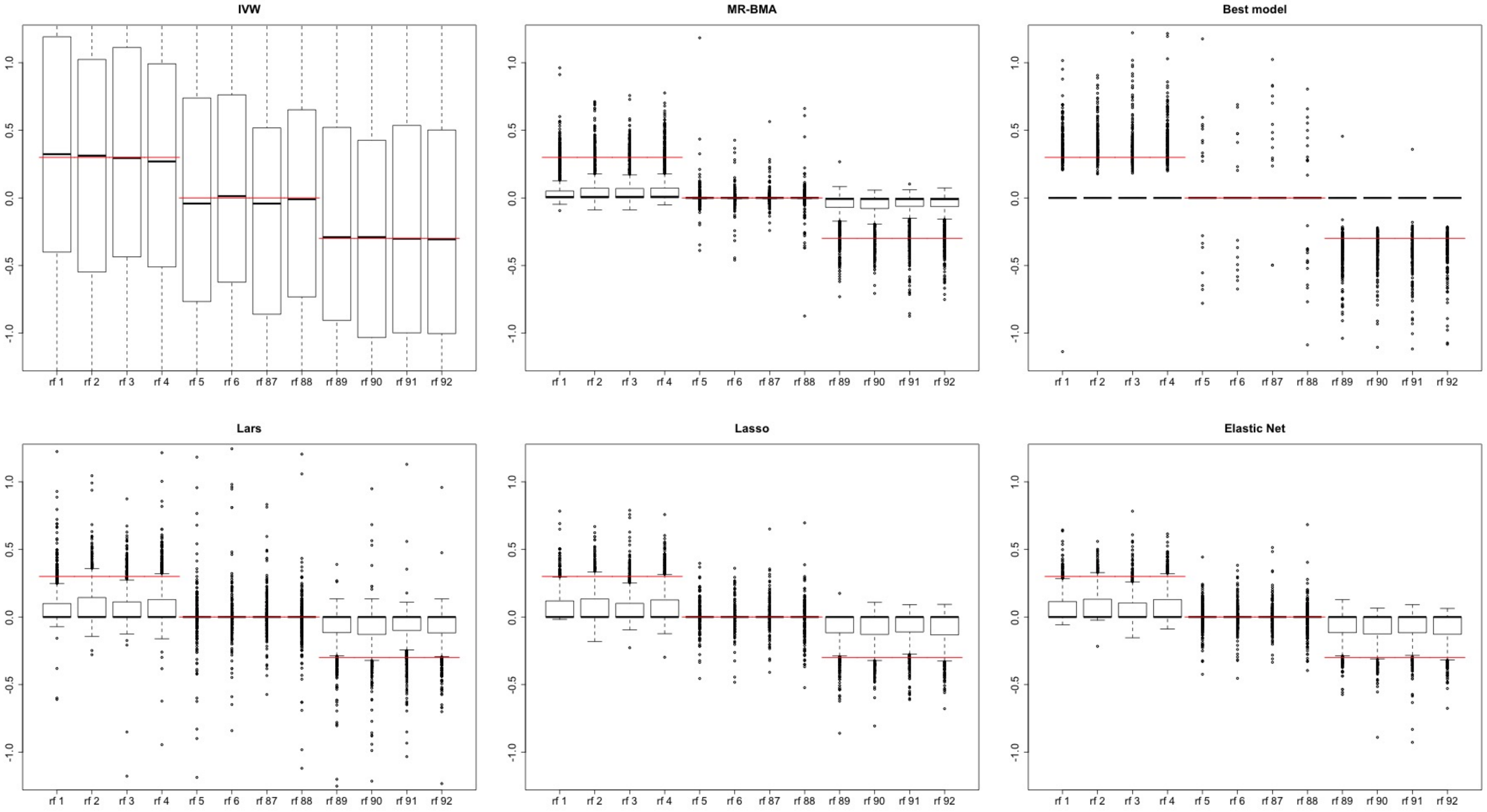

Figure 10: Boxplots of the causal effect estimates for setting B including a large number of risk factors ($d = 92$) of which the first four have a positive and the final 4 have a negative causal effect. Risk factors 7 to 86 are omitted. The true causal effects are marked in red. From top left to bottom right are the competing approaches: IVW, MR-BMA, best model, Lars, Lasso, and Elastic Net. Proportion of variance explained is set to 0.3.

#### 2.2 Supplementary Figures: Simulation results on blood cell trait data

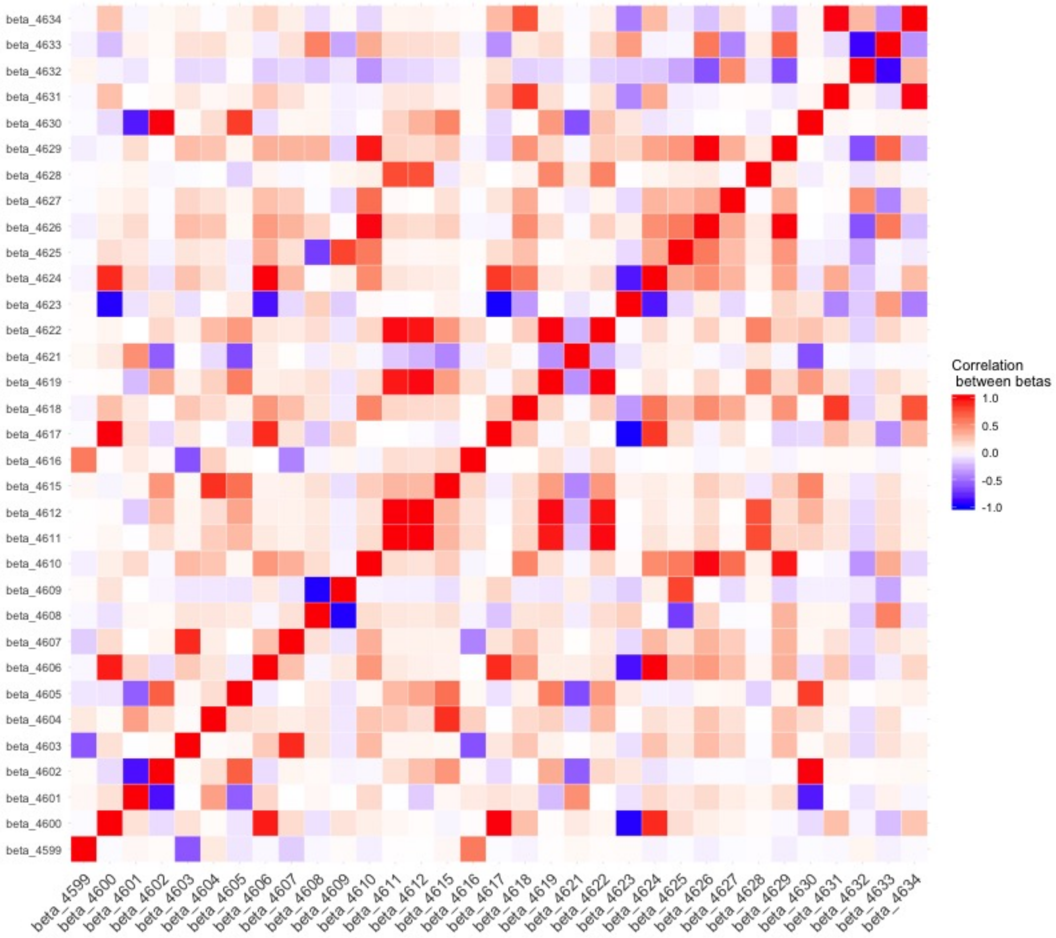

Figure 11: Genetic correlation between blood cell traits based on the $n = 2667$ genetic variants used as instrumental variables.

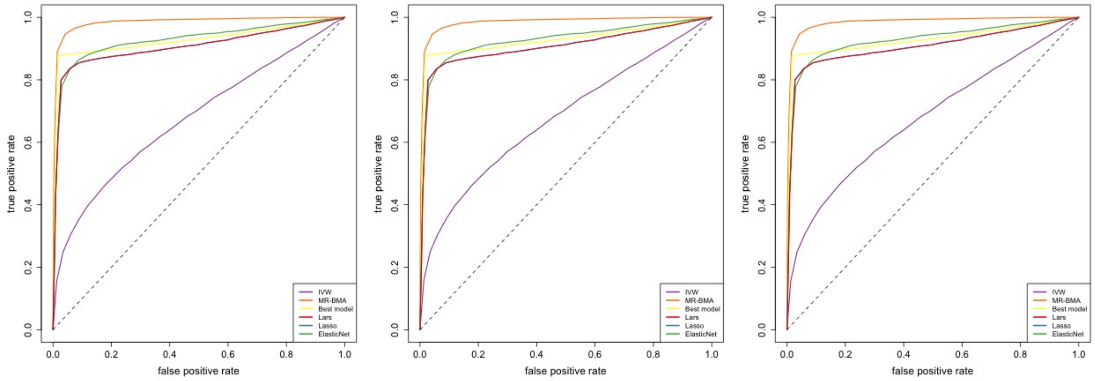

Figure 12: Receiver operating characteristic (ROC) curve for setting A including ($d = 33$) blood cell traits as risk factors of which four are true positive effects. Proportion of variance explained is set to 0.1 (left) 0.3 (middle) and 0.5 (right).

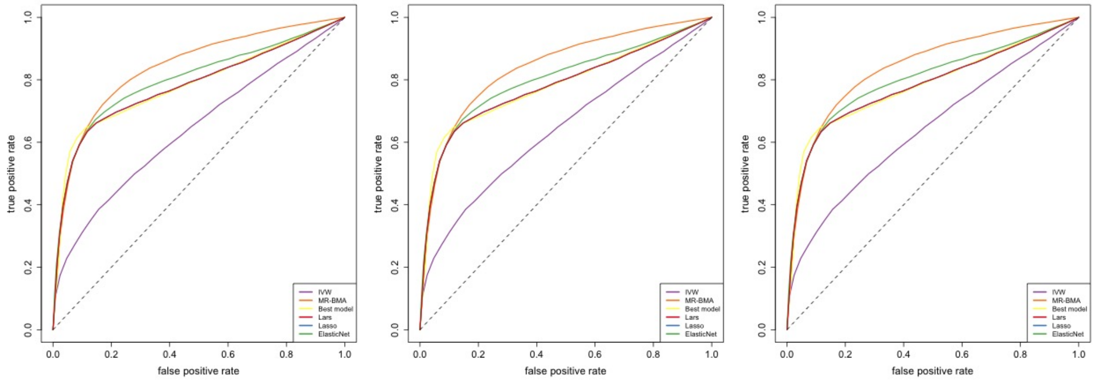

Figure 13: Receiver operating characteristic (ROC) curve for setting B including ($d = 33$) blood cell traits as risk factors of which four have true positive effect and another four have true negative effect. Proportion of variance explained is set to 0.1 (left) 0.3 (middle) and 0.5 (right).

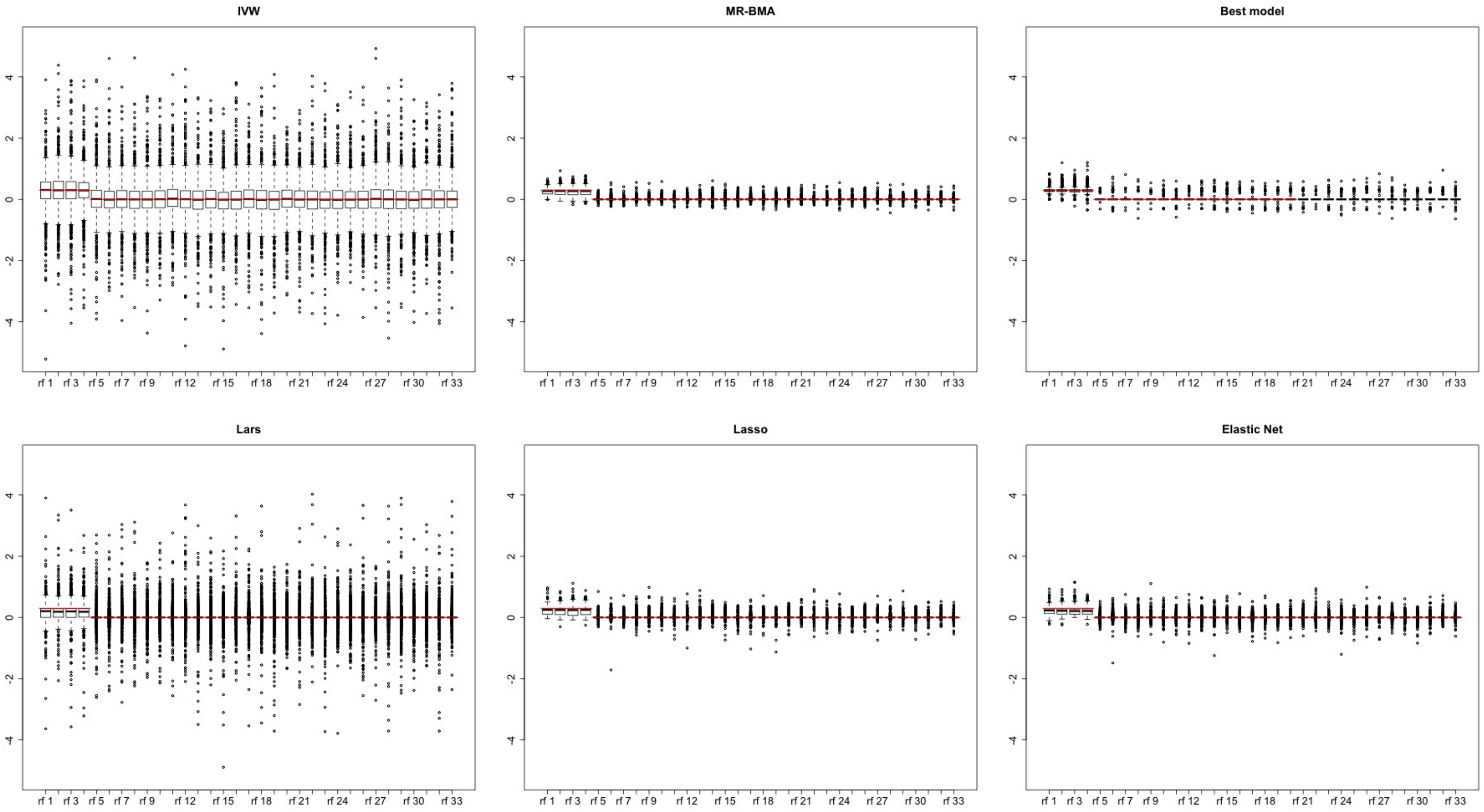

Figure 14: Boxplots of the causal effect estimates for setting A for the blood cell traits ($d = 33$), of which the first four are true positive effects. The true causal effects are marked in red. From top left to bottom right are the competing approaches: IVW, MR-BMA, best model and Lars, Lasso and Elastic Net (all tuned with cross-validation). Proportion of variance explained is set to 0.3.

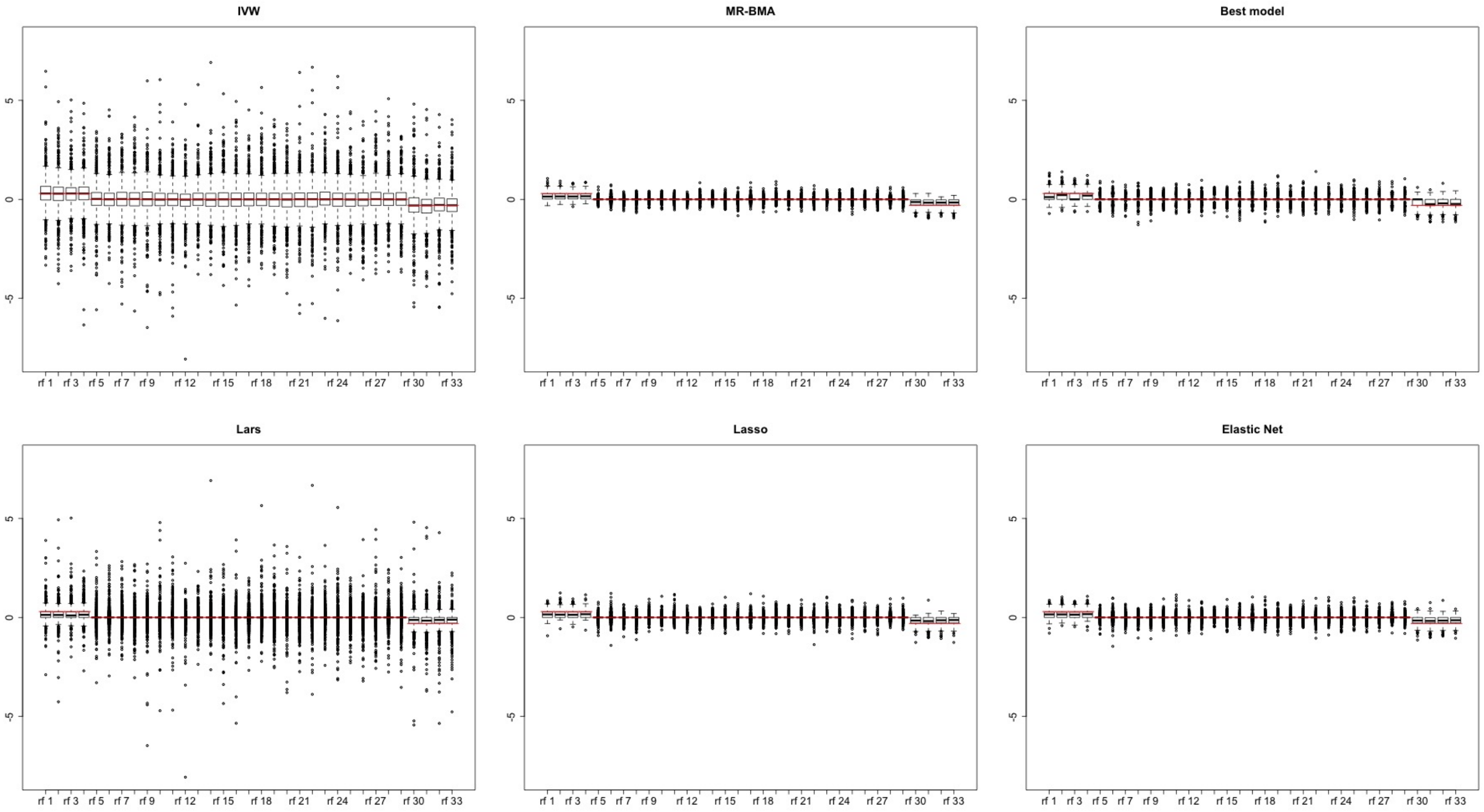

Figure 15: Boxplots of the causal effect estimates for setting B for the blood cell traits ($d = 33$), of which the first four have a positive and the last four have a negative causal effect. The true causal effects are marked in red. From top left to bottom right are the competing approaches: IVW, MR-BMA, best model and Lars, Lasso and Elastic Net (all tuned with cross-validation). Proportion of variance explained is set to 0.3.

### 3 Application: Metabolites as risk factors for age-related macular degeneration

#### 3.1 Supplementary Tables

A) Model averaging

| | risk factor | *MIP* | $\hat{\theta}_{\text{MACE}}$ |
|---|---|---|---|
| 1 | LDL.D | 0.527 | -0.229 |
| 2 | XS.VLDL.TG | 0.247 | -0.124 |
| 3 | S.HDL.TG | 0.236 | -0.101 |
| 4 | IDL.TG | 0.213 | -0.108 |
| 5 | XXL.VLDL.TG | 0.188 | 0.095 |
| 6 | S.VLDL.TG | 0.175 | -0.070 |
| 7 | S.LDL.C | 0.137 | 0.059 |
| 8 | Serum.TG | 0.137 | -0.062 |
| 9 | Est.C | 0.097 | 0.030 |
| 10 | XL.HDL.C | 0.085 | 0.021 |

B) Individual models

| | risk factor(s) | *PP* | $\hat{\theta}_{\gamma}$ |
|---|---|---|---|
| 1 | LDL.D,S.HDL.TG | 0.062 | -0.376,-0.398 |
| 2 | LDL.D,S.VLDL.TG | 0.052 | -0.485,-0.379 |
| 3 | LDL.D,Serum.TG | 0.020 | -0.454,-0.365 |
| 4 | S.HDL.TG | 0.019 | -0.433 |
| 5 | Est.C,IDL.TG | 0.019 | 0.393,-0.625 |
| 6 | LDL.D,XS.VLDL.TG | 0.018 | -0.339,-0.324 |
| 7 | XS.VLDL.TG | 0.017 | -0.373 |
| 8 | LDL.D,M.VLDL.TG | 0.014 | -0.545,-0.408 |
| 9 | S.HDL.TG,XXL.VLDL.TG | 0.013 | -0.653,0.45 |
| 10 | IDL.TG | 0.009 | -0.343 |

Table 1: Ranking of risk factors (top ten) for age-related macular degeneration according to their marginal inclusion probability (*MIP*) A) and the best ten individual model according to their posterior probability (*PP*) B). Calculation is based on all genetic variants $n = 148$ including the *LIPC* region. Abbreviations: *MIP*=marginal inclusion probability, MACE= model-averaged causal effect, *PP*=posterior probability for indivudal model.

| | rs | region | $q$ M1 | $q$ M2 | $q$ M3 | max $q$ |
|---|---|---|---|---|---|---|
| 1 | rs6859 | APOE | 17.007 | 17.388 | 17.132 | 17.388 |
| 2 | rs492602 | FUT2 | 15.526 | 13.899 | 14.591 | 15.526 |
| 3 | rs4465830 | ZNF335 | 7.395 | 11.127 | 14.223 | 14.223 |
| 4 | rs174532 | MYRF | 11.939 | 11.078 | 11.517 | 11.939 |
| 5 | rs6489818 | MAPKAPK5 | 11.226 | 10.857 | 10.68 | 11.226 |
| 6 | rs103294 | AC245884.7 | 8.857 | 9.255 | 9.504 | 9.504 |
| 7 | rs3817588 | GCKR | 7.263 | 8.095 | 8.411 | 8.411 |
| 8 | rs261342 | LIPC | 7.11 | 8.107 | 5.747 | 8.107 |
| 9 | rs903319 | SLC2A2 | 8.06 | 6.567 | 6.276 | 8.06 |
| 10 | rs2587534 | AL160408.6 | 6.498 | 6.063 | 6.999 | 6.999 |
| 11 | rs2710642 | EHBP1 | 6.662 | 6.955 | 6.538 | 6.955 |
| 12 | rs9491696 | RSPO3 | 6.317 | 5.658 | 5.966 | 6.317 |
| 13 | rs1689797 | LINC01344 | 4.638 | 5.325 | 6.079 | 6.079 |
| 14 | rs6882076 | TIMD4 | 5.742 | 4.023 | 3.706 | 5.742 |
| 15 | rs8176720 | ABO | 5.415 | 4.972 | 5.334 | 5.415 |
| 16 | rs688 | LDLR | 4.85 | 5.178 | 4.694 | 5.178 |
| 17 | rs1781930 | AKR1C8P | 4.978 | 4.585 | 4.445 | 4.978 |
| 18 | rs702485 | DAGLB | 4.863 | 3.892 | 4.335 | 4.863 |
| 19 | rs38855 | MET | 4.636 | 3.896 | 4.858 | 4.858 |
| 20 | rs2925979 | CMIP | 4.66 | 4.516 | 4.243 | 4.66 |
| 21 | rs7703051 | HMGCR | 4.581 | 3.988 | 3.928 | 4.581 |
| 22 | rs2602836 | ADH5 | 3.724 | 4.357 | 4.528 | 4.528 |
| 23 | rs3741414 | INHBC | 3.873 | 4.434 | 4.158 | 4.434 |
| 24 | rs4148218 | ABCG8 | 3.967 | 3.592 | 3.666 | 3.967 |
| 25 | rs3822072 | FAM13A | 3.549 | 3.858 | 3.811 | 3.858 |
| 26 | rs5880 | CETP | 1.127 | 2.123 | 3.679 | 3.679 |
| 27 | rs6680658 | GALNT2 | 3.124 | 3.675 | 3.457 | 3.675 |
| 28 | rs9930333 | FTO | 3.351 | 3.428 | 3.04 | 3.428 |
| 29 | rs7225700 | THCAT158 | 3.127 | 3.305 | 3.381 | 3.381 |
| 30 | rs217386 | NPC1L1 | 1.959 | 3.311 | 2.665 | 3.311 |

Table 2: $q$-statistic for all $n = 148$ genetic variants for the best individual model 1 (M1: LDL.D and S.HDL.TG), model 2 (M2: LDL.D and S.VLDL.TG), and model 3 (M3: LDL.D and Serum.TG) and the maximum $q$ of each variant in all models used for diagnostics. This table displays the 30 variants with the largest maximum $q$ and the region they fall in.

| | rs | region | $Cd$ M1 | $Cd$ M2 | $Cd$ M3 | max $Cd$ |
|---|---|---|---|---|---|---|
| 1 | rs261342 | LIPC | **0.989** | **1.087** | **0.871** | 1.087 |
| 2 | rs4465830 | ZNF335 | 0.188 | 0.108 | 0.056 | 0.188 |
| 3 | rs3817588 | GCKR | 0.058 | 0.085 | 0.105 | 0.105 |
| 4 | rs6859 | APOE | 0.081 | 0.076 | 0.087 | 0.087 |
| 5 | rs5880 | CETP | 0.056 | 0.071 | 0.081 | 0.081 |
| 6 | rs174532 | MYRF | 0.062 | 0.062 | 0.061 | 0.062 |
| 7 | rs686030 | TTC39B | 0.054 | 0.04 | 0.052 | 0.054 |
| 8 | rs7703051 | HMGCR | 0.039 | 0.045 | 0.05 | 0.05 |
| 9 | rs103294 | AC245884.7 | 0.045 | 0.044 | 0.044 | 0.045 |
| 10 | rs10401969 | SUGP1 | 0.009 | 0.025 | 0.043 | 0.043 |
| 11 | rs1689797 | LINC01344 | 0.037 | 0.031 | 0.026 | 0.037 |
| 12 | rs2710642 | EHBP1 | 0.031 | 0.033 | 0.03 | 0.033 |
| 13 | rs2587534 | AL160408.6 | 0.02 | 0.018 | 0.024 | 0.024 |
| 14 | rs10493326 | DOCK7 | 0.011 | 0.017 | 0.023 | 0.023 |
| 15 | rs894210 | intergenic | 0.015 | 0.022 | 0.02 | 0.022 |
| 16 | rs6882076 | TIMD4 | 0.006 | 0.016 | 0.02 | 0.02 |
| 17 | rs903319 | SLC2A2 | 0.02 | 0.008 | 0.007 | 0.02 |
| 18 | rs515135 | APOB(intergenic) | 0.019 | 0.011 | 0.012 | 0.019 |
| 19 | rs799160 | intergenic | 0.017 | 0.016 | 0.019 | 0.019 |
| 20 | rs3741414 | INHBC | 0.01 | 0.016 | 0.013 | 0.016 |
| 21 | rs1515110 | NR | 0.014 | 0.01 | 0.007 | 0.014 |
| 22 | rs1800562 | HFE | 0.01 | 0.012 | 0.012 | 0.012 |
| 23 | rs2068888 | CYP26A1 | 0.012 | 0.011 | 0.011 | 0.012 |
| 24 | rs7225700 | THCAT158 | 0.011 | 0.012 | 0.012 | 0.012 |
| 25 | rs492602 | FUT2 | 0.011 | 0.002 | 0.005 | 0.011 |
| 26 | rs38855 | MET | 0.008 | 0.003 | 0.01 | 0.01 |
| 27 | rs688 | LDLR | 0.007 | 0.01 | 0.006 | 0.01 |
| 28 | rs6680658 | GALNT2 | 0.005 | 0.01 | 0.009 | 0.01 |
| 29 | rs3198697 | PDXDC1 | 0.007 | 0.01 | 0.01 | 0.01 |
| 30 | rs2326077 | intergenic | 0.006 | 0.006 | 0.01 | 0.01 |
| | threshold | | 0.696 | 0.696 | 0.696 | |

Table 3: Cook's distance ($Cd$) based on $n = 148$ genetic variants including *LIPC* for the best individual model 1 (M1: LDL.D and S.HDL.TG), model 2 (M2: LDL.D and S.VLDL.TG), and model 3 (M3: LDL.D and Serum.TG). The final line gives the suggested cut-off for Cook's distance and variants with $Cd$ above this threshold are given in bold. This table displays the 30 variants with the largest maximum Cook's distance and the region they fall in.

| | risk factor | *MIP* | $\hat{\theta}_{\text{MACE}}$ |
|---|---|---|---|
| 1 | XL.HDL.C | 0.700 | 0.344 |
| 2 | L.HDL.C | 0.229 | 0.087 |
| 3 | HDL.D | 0.087 | 0.022 |
| 4 | XS.VLDL.TG | 0.082 | -0.019 |
| 5 | LDL.D | 0.074 | -0.018 |
| 6 | IDL.TG | 0.066 | -0.012 |
| 7 | XXL.VLDL.TG | 0.063 | 0.018 |
| 8 | S.VLDL.TG | 0.062 | -0.014 |
| 9 | Serum.TG | 0.061 | -0.014 |
| 10 | Serum.C | 0.054 | -0.011 |
| 11 | HDL.C | 0.051 | 0.009 |
| 12 | M.HDL.C | 0.048 | -0.010 |
| 13 | S.HDL.TG | 0.047 | -0.006 |
| 14 | XL.HDL.TG | 0.045 | 0.005 |
| 15 | M.VLDL.C | 0.043 | -0.005 |
| 16 | S.VLDL.C | 0.043 | -0.005 |
| 17 | ApoA1 | 0.040 | -0.007 |
| 18 | M.VLDL.TG | 0.039 | 0.006 |
| 19 | ApoB | 0.038 | -0.004 |
| 20 | L.VLDL.C | 0.038 | -0.005 |
| 21 | XL.VLDL.TG | 0.034 | -0.003 |
| 22 | L.VLDL.TG | 0.033 | -0.001 |
| 23 | S.LDL.C | 0.033 | 0.001 |
| 24 | LDL.C | 0.031 | -0.003 |
| 25 | IDL.C | 0.029 | -0.001 |
| 26 | SM | 0.027 | -0.003 |
| 27 | VLDL.D | 0.027 | 0.002 |
| 28 | Tot.FA | 0.026 | -0.001 |
| 29 | Est.C | 0.026 | 0.001 |
| 30 | TotPG | 0.026 | -0.002 |

Table 4: Ranking of risk factors for age-related macular degeneration according to their marginal inclusion probability (*MIP*) after excluding genetic variants in the *LIPC, FUT2* and *APOE* region $n = 145$. Abbreviations: *MIP*=marginal inclusion probability, MACE= model-averaged causal effect.

| | rs | region | Q M1 | Q M2 | Q M3 | Q M4 | Q M5 | max Q |
|---|---|---|---|---|---|---|---|---|
| 1 | rs103294 | AC245884.7 | 13.03 | 13.155 | 11.936 | 11.203 | 14.449 | 14.449 |
| 2 | rs6489818 | MAPKAPK5 | 11.244 | 9.575 | 10.53 | 10.356 | 9.883 | 11.244 |
| 3 | rs6882076 | TIMD4 | 9.536 | 9.118 | 6.708 | 6.503 | 10.504 | 10.504 |
| 4 | rs2587534 | AL160408.6 | 5.931 | 8.936 | 6.551 | 6.735 | 8.409 | 8.936 |
| 5 | rs903319 | SLC2A2 | 7.514 | 6.651 | 7.275 | 7.255 | 6.379 | 7.514 |
| 6 | rs3817588 | GCKR | 4.698 | 6.3 | 7.015 | 6.495 | 7.051 | 7.051 |
| 7 | rs1689797 | LINC01344 | 6.403 | 4.747 | 4.635 | 4.587 | 5.648 | 6.403 |
| 8 | rs8176720 | ABO | 3.929 | 6.312 | 4.592 | 4.734 | 5.197 | 6.312 |
| 9 | rs38855 | MET | 3.768 | 5.98 | 5.082 | 4.973 | 5.205 | 5.98 |
| 10 | rs9491696 | RSPO3 | 5.651 | 5.974 | 5.017 | 4.971 | 5.479 | 5.974 |
| 11 | rs7703051 | HMGCR | 5.974 | 3.24 | 3.246 | 3.319 | 4.009 | 5.974 |
| 12 | rs688 | LDLR | 2.562 | 5.557 | 3.97 | 4.071 | 4.856 | 5.557 |
| 13 | rs5880 | CETP | 5.433 | 2.877 | 2.687 | 2.73 | 4.246 | 5.433 |
| 14 | rs1781930 | AKR1C8P | 5.176 | 4.259 | 4.996 | 5.072 | 4.851 | 5.176 |
| 15 | rs3822072 | FAM13A | 5.105 | 3.376 | 4.504 | 4.606 | 4.099 | 5.105 |
| 16 | rs2923084 | AMPD3 | 5.067 | 2.814 | 2.956 | 2.944 | 3.933 | 5.067 |
| 17 | rs9693857 | AC022784.6 | 4.752 | 3.147 | 3.966 | 4.246 | 3.601 | 4.752 |
| 18 | rs2710642 | EHBP1 | 3.632 | 3.432 | 4.381 | 4.714 | 3.318 | 4.714 |
| 19 | rs174532 | MYRF | 2.708 | 4.701 | 3.405 | 3.927 | 4.12 | 4.701 |
| 20 | rs6680658 | GALNT2 | 3.216 | 3.885 | 3.926 | 3.527 | 3.577 | 3.926 |
| 21 | rs686030 | TTC39B | 1.58 | 3.558 | 1.7 | 1.393 | 3.913 | 3.913 |
| 22 | rs702485 | DAGLB | 3.569 | 3.439 | 3.887 | 3.768 | 3.597 | 3.887 |
| 23 | rs9930333 | FTO | 3.872 | 2.154 | 3.299 | 3.245 | 2.299 | 3.872 |
| 24 | rs17789218 | intergenic | 3.72 | 2.12 | 3.145 | 3.219 | 3.512 | 3.72 |
| 25 | rs2068888 | CYP26A1 | 3.714 | 1.944 | 2.47 | 2.627 | 2.291 | 3.714 |
| 26 | rs9686661 | C5orf67 | 3.702 | 1.258 | 2.31 | 2.597 | 1.811 | 3.702 |
| 27 | rs2297374 | SLC22A1 | 3.294 | 2.614 | 2.716 | 2.554 | 3.608 | 3.608 |
| 28 | rs2925979 | CMIP | 3.135 | 3.14 | 3.417 | 3.486 | 3.142 | 3.486 |
| 29 | rs3741414 | INHBC | 2.203 | 2.149 | 3.335 | 3.438 | 1.8 | 3.438 |
| 30 | rs7264396 | FER1L4 | 2.74 | 3.251 | 2.562 | 2.372 | 3.438 | 3.438 |

Table 5: $q$-statistic for $n = 145$ genetic variants after excluding *LIPC, FUT2* and *APOE* the best individual model 1 (M1: XL.HDL.C), model 2 (M2: L.HDL.C), model 3 (M3: XL.HDL.C and XS.VLDL.TG), model 4 (M4: IDL.TG and XL.HDL.C), model 5 (M5: HDL.D), and the maximum $q$ of each variant in all models used for diagnostics. This table displays the 30 variants with the largest maximum $q$ and the region they fall in.

| | rs | region | $Cd$ M1 | $Cd$ M2 | $Cd$ M3 | $Cd$ M4 | $Cd$ M5 | max $Cd$ |
|---|---|---|---|---|---|---|---|---|
| 1 | rs4465830 | ZNF335 | 0.216 | 0.311 | 0.106 | 0.113 | 0.271 | 0.311 |
| 2 | rs5880 | CETP | 0.234 | 0.277 | 0.122 | 0.122 | 0.297 | 0.297 |
| 3 | rs1689797 | LINC01344 | 0.061 | 0.098 | 0.047 | 0.048 | 0.086 | 0.098 |
| 4 | rs686030 | TTC39B | 0.072 | 0.062 | 0.04 | 0.033 | 0.062 | 0.072 |
| 5 | rs6882076 | TIMD4 | 0.004 | 0.001 | 0.061 | 0.07 | 0.016 | 0.07 |
| 6 | rs3817588 | GCKR | 0.005 | 0 | 0.062 | 0.037 | 0.005 | 0.062 |
| 7 | rs174532 | MYRF | 0.052 | 0.027 | 0.039 | 0.057 | 0.039 | 0.057 |
| 8 | rs13107325 | SLC39A8 | 0.001 | 0.032 | 0.001 | 0 | 0.056 | 0.056 |
| 9 | rs7703051 | HMGCR | 0.001 | 0.027 | 0.05 | 0.048 | 0.019 | 0.05 |
| 10 | rs903319 | SLC2A2 | 0.047 | 0.025 | 0.024 | 0.024 | 0.021 | 0.047 |
| 11 | rs894210 | intergenic | 0.008 | 0.046 | 0.023 | 0.011 | 0.018 | 0.046 |
| 12 | rs10773105 | SCARB1 | 0.015 | 0.042 | 0.009 | 0.008 | 0.04 | 0.042 |
| 13 | rs103294 | AC245884.7 | 0.028 | 0.026 | 0.023 | 0.039 | 0.009 | 0.039 |
| 14 | rs998584 | VEGFA(intergenic) | 0 | 0.039 | 0.005 | 0.002 | 0.013 | 0.039 |
| 15 | rs17789218 | intergenic | 0.034 | 0.003 | 0.017 | 0.017 | 0.031 | 0.034 |
| 16 | rs1800961 | HNF4A | 0.015 | 0.033 | 0.013 | 0.014 | 0.011 | 0.033 |
| 17 | rs2923084 | AMPD3 | 0.001 | 0.009 | 0.03 | 0.031 | 0.002 | 0.031 |
| 18 | rs688 | LDLR | 0.005 | 0.011 | 0.025 | 0.029 | 0.004 | 0.029 |
| 19 | rs2587534 | AL160408.6 | 0.026 | 0 | 0.018 | 0.021 | 0.001 | 0.026 |
| 20 | rs1515110 | NR | 0.008 | 0.025 | 0.012 | 0.008 | 0.016 | 0.025 |
| 21 | rs7897379 | REEP3 | 0.017 | 0.018 | 0.013 | 0.009 | 0.024 | 0.024 |
| 22 | rs10493326 | DOCK7 | 0.003 | 0.017 | 0.021 | 0.014 | 0.004 | 0.021 |
| 23 | rs499974 | RN7SL786P | 0.016 | 0.021 | 0.009 | 0.009 | 0.018 | 0.021 |
| 24 | rs9491696 | RSPO3 | 0.016 | 0.013 | 0.01 | 0.011 | 0.02 | 0.02 |
| 25 | rs3741414 | INHBC | 0.003 | 0.002 | 0.016 | 0.019 | 0.001 | 0.019 |
| 26 | rs9686661 | C5orf67 | 0.013 | 0.002 | 0.017 | 0.014 | 0 | 0.017 |
| 27 | rs38855 | MET | 0 | 0.014 | 0.017 | 0.015 | 0.005 | 0.017 |
| 28 | rs2602836 | ADH5 | 0.016 | 0.011 | 0.011 | 0.011 | 0.016 | 0.016 |
| 29 | rs2278236 | ANGPTL4 | 0.01 | 0.015 | 0.005 | 0.005 | 0.01 | 0.015 |
| 30 | rs702485 | DAGLB | 0.013 | 0.014 | 0.008 | 0.007 | 0.014 | 0.014 |
| | | | 0.457 | 0.457 | 0.697 | 0.697 | 0.457 | |

Table 6: Cook's distance ($Cd$) based on $n = 145$ genetic variants after excluding *LIPC, FUT2* and *APOE* the best individual model 1 (M1: XL.HDL.C), model 2 (M2: L.HDL.C), model 3 (M3: XL.HDL.C and XS.VLDL.TG), model 4 (M4: IDL.TG and XL.HDL.C), model 5 (M5: HDL.D), the final line gives the suggested cut-off for Cook's distance and this time, there are no variants with $Cd$ above this threshold. This table displays the 30 variants with the largest maximum Cook's distance and the region they fall in.

$p = 0.01$

| # | risk factor | *MIP* | $\hat{\theta}_{\mathrm{MACE}}$ |
|---|---|---|---|
| 1 | XL.HDL.C | 0.608 | 0.308 |
| 2 | L.HDL.C | 0.283 | 0.109 |
| 3 | HDL.D | 0.087 | 0.030 |
| 4 | HDL.C | 0.024 | 0.008 |
| 5 | XS.VLDL.TG | 0.011 | -0.002 |
| 6 | IDL.TG | 0.009 | -0.002 |
| 7 | S.HDL.TG | 0.009 | -0.002 |
| 8 | LDL.D | 0.007 | -0.002 |
| 9 | Serum.C | 0.007 | -0.001 |
| 10 | S.VLDL.TG | 0.007 | -0.001 |

$p = 0.05$

| # | risk factor | *MIP* | $\hat{\theta}_{\mathrm{MACE}}$ |
|---|---|---|---|
| 1 | XL.HDL.C | 0.663 | 0.330 |
| 2 | L.HDL.C | 0.249 | 0.095 |
| 3 | HDL.D | 0.084 | 0.026 |
| 4 | XS.VLDL.TG | 0.047 | -0.010 |
| 5 | IDL.TG | 0.040 | -0.007 |
| 6 | LDL.D | 0.037 | -0.008 |
| 7 | HDL.C | 0.035 | 0.008 |
| 8 | S.VLDL.TG | 0.032 | -0.006 |
| 9 | Serum.C | 0.030 | -0.005 |
| 10 | Serum.TG | 0.029 | -0.006 |

$p = 0.1$

| # | risk factor | *MIP* | $\hat{\theta}_{\mathrm{MACE}}$ |
|---|---|---|---|
| 1 | XL.HDL.C | 0.70 | 0.34 |
| 2 | L.HDL.C | 0.23 | 0.09 |
| 3 | HDL.D | 0.09 | 0.02 |
| 4 | XS.VLDL.TG | 0.08 | -0.02 |
| 5 | LDL.D | 0.07 | -0.02 |
| 6 | IDL.TG | 0.07 | -0.01 |
| 7 | S.VLDL.TG | 0.06 | -0.01 |
| 8 | XXL.VLDL.TG | 0.06 | 0.02 |
| 9 | Serum.TG | 0.06 | -0.01 |
| 10 | Serum.C | 0.05 | -0.01 |

$p = 0.2$

| # | risk factor | *MIP* | $\hat{\theta}_{\mathrm{MACE}}$ |
|---|---|---|---|
| 1 | XL.HDL.C | 0.700 | 0.344 |
| 2 | L.HDL.C | 0.229 | 0.087 |
| 3 | HDL.D | 0.087 | 0.022 |
| 4 | XS.VLDL.TG | 0.082 | -0.019 |
| 5 | LDL.D | 0.075 | -0.018 |
| 6 | IDL.TG | 0.067 | -0.013 |
| 7 | S.VLDL.TG | 0.062 | -0.014 |
| 8 | XXL.VLDL.TG | 0.061 | 0.018 |
| 9 | Serum.TG | 0.061 | -0.014 |
| 10 | Serum.C | 0.053 | -0.010 |

$p = 0.3$

| # | risk factor | *MIP* | $\hat{\theta}_{\mathrm{MACE}}$ |
|---|---|---|---|
| 1 | XL.HDL.C | 0.675 | 0.315 |
| 2 | L.HDL.C | 0.302 | 0.126 |
| 3 | XXL.VLDL.TG | 0.300 | 0.121 |
| 4 | LDL.D | 0.244 | -0.073 |
| 5 | Serum.TG | 0.212 | -0.065 |
| 6 | XS.VLDL.TG | 0.197 | -0.052 |
| 7 | S.VLDL.TG | 0.190 | -0.053 |
| 8 | M.VLDL.TG | 0.173 | 0.048 |
| 9 | Serum.C | 0.165 | -0.053 |
| 10 | ApoA1 | 0.152 | -0.038 |

Table 7: Parameter check for the prior probability $p$, ranging from $p = 0.01$ to $p = 0.3$. This reflects 0.3 to 9 expected causal risk factors. We used $p = 0.1$ reflecting an a priori expected number of 3 causal risk factors in the main analysis. Abbreviations: *MIP*=marginal inclusion probability, MACE= model-averaged causal effect.

$\sigma = 0.1$

| # | risk factor | *MIP* | $\hat{\theta}_{\text{MACE}}$ |
|---|---|---|---|
| 1 | XL.HDL.C | 0.52 | 0.13 |
| 2 | L.HDL.C | 0.42 | 0.09 |
| 3 | HDL.D | 0.27 | 0.05 |
| 4 | LDL.D | 0.15 | -0.02 |
| 5 | HDL.C | 0.14 | 0.02 |
| 6 | XS.VLDL.TG | 0.13 | -0.02 |
| 7 | S.HDL.TG | 0.13 | -0.02 |
| 8 | S.VLDL.TG | 0.11 | -0.01 |
| 9 | IDL.TG | 0.10 | -0.01 |
| 10 | Serum.TG | 0.09 | -0.01 |

$\sigma = 0.3$

| # | risk factor | *MIP* | $\hat{\theta}_{\text{MACE}}$ |
|---|---|---|---|
| 1 | XL.HDL.C | 0.69 | 0.32 |
| 2 | L.HDL.C | 0.25 | 0.09 |
| 3 | XS.VLDL.TG | 0.11 | -0.02 |
| 4 | HDL.D | 0.11 | 0.03 |
| 5 | LDL.D | 0.10 | -0.02 |
| 6 | IDL.TG | 0.08 | -0.01 |
| 7 | S.VLDL.TG | 0.08 | -0.02 |
| 8 | XXL.VLDL.TG | 0.08 | 0.02 |
| 9 | Serum.TG | 0.07 | -0.01 |
| 10 | S.HDL.TG | 0.06 | -0.01 |

$\sigma = 0.5$

| # | risk factor | *MIP* | $\hat{\theta}_{\text{MACE}}$ |
|---|---|---|---|
| 1 | XL.HDL.C | 0.70 | 0.34 |
| 2 | L.HDL.C | 0.23 | 0.09 |
| 3 | HDL.D | 0.09 | 0.02 |
| 4 | XS.VLDL.TG | 0.08 | -0.02 |
| 5 | LDL.D | 0.07 | -0.02 |
| 6 | IDL.TG | 0.07 | -0.01 |
| 7 | S.VLDL.TG | 0.06 | -0.01 |
| 8 | XXL.VLDL.TG | 0.06 | 0.02 |
| 9 | Serum.TG | 0.06 | -0.01 |
| 10 | Serum.C | 0.05 | -0.01 |

$\sigma = 0.7$

| # | risk factor | *MIP* | $\hat{\theta}_{\text{MACE}}$ |
|---|---|---|---|
| 1 | XL.HDL.C | 0.69 | 0.35 |
| 2 | L.HDL.C | 0.23 | 0.09 |
| 3 | HDL.D | 0.08 | 0.02 |
| 4 | XS.VLDL.TG | 0.07 | -0.02 |
| 5 | LDL.D | 0.06 | -0.01 |
| 6 | IDL.TG | 0.05 | -0.01 |
| 7 | S.VLDL.TG | 0.05 | -0.01 |
| 8 | Serum.TG | 0.05 | -0.01 |
| 9 | XXL.VLDL.TG | 0.05 | 0.02 |
| 10 | Serum.C | 0.05 | -0.01 |

Table 8: Parameter check for the prior variance $\sigma^2$, ranging from $\sigma = 0.1$ to $\sigma = 0.7$. We used $\sigma = 0.5$ in the main analysis. Abbreviations: *MIP*=marginal inclusion probability, MACE= model-averaged causal effect.

| | risk factor | beta | $p-$value |
|---|---|---|---|
| 1 | Serum.C | -2.033 | 0.004 |
| 2 | LDL.C | -1.808 | 0.014 |
| 3 | IDL.C | 2.156 | 0.014 |
| 4 | XXL.VLDL.TG | 1.075 | 0.015 |
| 5 | M.VLDL.TG | 1.769 | 0.019 |
| 6 | LDL.D | -0.937 | 0.032 |
| 7 | S.LDL.C | 1.302 | 0.064 |
| 8 | S.VLDL.C | 1.046 | 0.066 |
| 9 | L.HDL.C | 1.350 | 0.129 |
| 10 | S.HDL.TG | 0.562 | 0.175 |
| 11 | SM | -0.221 | 0.223 |
| 12 | VLDL.D | -0.497 | 0.250 |
| 13 | ApoA1 | -0.390 | 0.318 |
| 14 | XS.VLDL.TG | -1.015 | 0.330 |
| 15 | M.VLDL.C | -0.856 | 0.339 |
| 16 | Tot.FA | 0.350 | 0.359 |
| 17 | L.VLDL.TG | -0.616 | 0.371 |
| 18 | TotPG | -0.246 | 0.470 |
| 19 | Serum.TG | -0.771 | 0.525 |
| 20 | XL.VLDL.TG | -0.302 | 0.605 |
| 21 | IDL.TG | 0.398 | 0.654 |
| 22 | ApoB | 0.273 | 0.658 |
| 23 | L.VLDL.C | -0.241 | 0.670 |
| 24 | M.HDL.C | 0.098 | 0.814 |
| 25 | Est.C | 0.082 | 0.828 |
| 26 | HDL.C | -0.193 | 0.838 |
| 27 | XL.HDL.TG | 0.083 | 0.850 |
| 28 | XL.HDL.C | 0.079 | 0.868 |
| 29 | S.VLDL.TG | 0.066 | 0.932 |
| 30 | HDL.D | -0.029 | 0.958 |

Table 9: Ranking of risk factors for age-related macular degeneration using inverse-variance weighted (IVW) regression according to their $p-$value after excluding genetic variants in the *LIPC, FUT2* and *APOE* region $n = 145$. Abbreviations: beta=causal effect, p=$p$-value of the causal effect (not adjusted for multiple testing).

| | risk factor | beta L1 |
|---|---|---|
| 1 | L.HDL.C | 0.357 |
| 2 | LDL.D | -0.255 |
| 3 | XXL.VLDL.TG | 0.251 |
| 4 | S.VLDL.TG | -0.170 |
| 5 | M.HDL.C | -0.157 |
| 6 | XL.HDL.C | 0.115 |
| 7 | XL.VLDL.TG | -0.104 |
| 8 | ApoA1 | -0.093 |
| 9 | Est.C | 0.062 |
| 10 | Serum.TG | -0.010 |
| 11 | SM | -0.005 |
| | ApoB | 0 |
| | HDL.C | 0 |
| | HDL.D | 0 |
| | IDL.C | 0 |
| | IDL.TG | 0 |
| | L.VLDL.C | 0 |
| | L.VLDL.TG | 0 |
| | LDL.C | 0 |
| | M.VLDL.C | 0 |
| | M.VLDL.TG | 0 |
| | S.HDL.TG | 0 |
| | S.LDL.C | 0 |
| | S.VLDL.C | 0 |
| | Serum.C | 0 |
| | Tot.FA | 0 |
| | TotPG | 0 |
| | VLDL.D | 0 |
| | XL.HDL.TG | 0 |
| | XS.VLDL.TG | 0 |

Table 10: Ranking of risk factors for age-related macular degeneration using Lars regression according to their L1 regularised causal effect estimate after excluding genetic variants in the *LIPC, FUT2* and *APOE* region $n = 145$. Abbreviations: beta L1=L1 regularised causal effect.

| | risk factor | beta L1 |
|---|---|---|
| 1 | XL.HDL.C | 0.306 |
| 2 | XS.VLDL.TG | -0.102 |
| 3 | L.HDL.C | 0.092 |
| 4 | LDL.D | -0.039 |
| | ApoA1 | 0 |
| | ApoB | 0 |
| | Est.C | 0 |
| | HDL.C | 0 |
| | HDL.D | 0 |
| | IDL.C | 0 |
| | IDL.TG | 0 |
| | L.VLDL.C | 0 |
| | L.VLDL.TG | 0 |
| | LDL.C | 0 |
| | M.HDL.C | 0 |
| | M.VLDL.C | 0 |
| | M.VLDL.TG | 0 |
| | S.HDL.TG | 0 |
| | S.LDL.C | 0 |
| | S.VLDL.C | 0 |
| | S.VLDL.TG | 0 |
| | Serum.C | 0 |
| | Serum.TG | 0 |
| | SM | 0 |
| | Tot.FA | 0 |
| | TotPG | 0 |
| | VLDL.D | 0 |
| | XL.HDL.TG | 0 |
| | XL.VLDL.TG | 0 |
| | XXL.VLDL.TG | 0 |

Table 11: Ranking of risk factors for age-related macular degeneration using Lasso regression (L1 penalty) according to their regularised causal effect estimate after excluding genetic variants in the *LIPC, FUT2* and *APOE* region $n = 145$. Abbreviations: beta L1=L1 regularised causal effect.

| | risk factor | beta L1+L2 |
|---|---|---|
| 1 | L.HDL.C | 0.269 |
| 2 | XL.HDL.C | 0.176 |
| 3 | LDL.D | -0.172 |
| 4 | M.HDL.C | -0.137 |
| 5 | XXL.VLDL.TG | 0.117 |
| 6 | XS.VLDL.TG | -0.102 |
| 7 | S.VLDL.TG | -0.090 |
| 8 | Est.C | 0.065 |
| 9 | ApoA1 | -0.052 |
| 10 | Serum.C | -0.010 |
| | ApoB | 0 |
| | HDL.C | 0 |
| | HDL.D | 0 |
| | IDL.C | 0 |
| | IDL.TG | 0 |
| | L.VLDL.C | 0 |
| | L.VLDL.TG | 0 |
| | LDL.C | 0 |
| | M.VLDL.C | 0 |
| | M.VLDL.TG | 0 |
| | S.HDL.TG | 0 |
| | S.LDL.C | 0 |
| | S.VLDL.C | 0 |
| | Serum.TG | 0 |
| | SM | 0 |
| | Tot.FA | 0 |
| | TotPG | 0 |
| | VLDL.D | 0 |
| | XL.HDL.TG | 0 |
| | XL.VLDL.TG | 0 |

Table 12: Ranking of risk factors for age-related macular degeneration using Elastic Net regression (L1+L2 penalty) according to their regularised causal effect estimate after excluding genetic variants in the *LIPC, FUT2* and *APOE* region $n = 145$. Abbreviations: beta L1+L2=L1 and L2 regularised causal effect.

#### 3.2 Supplementary Figures

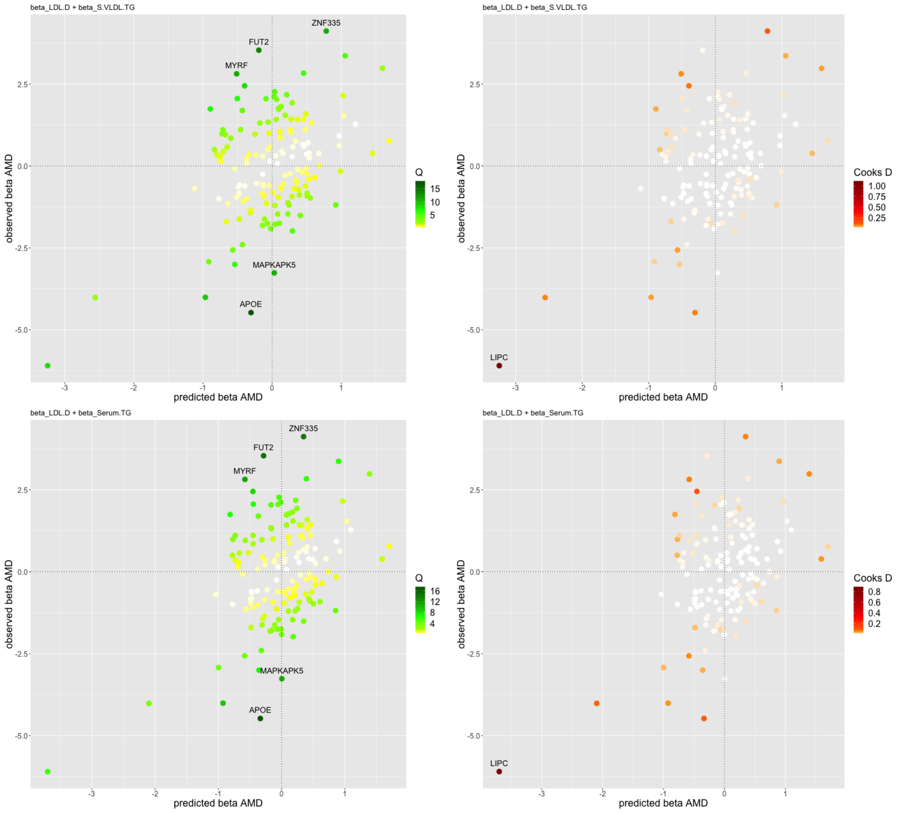

Figure 16: Diagnostic plots of the predicted associations with AMD ($x$-axis) based on model 2 (M2: LDL diameter (LDL.D) and TG in small VLDL (S.VLDL.TG)), model 3 (M3: LDL.D and Serum.TG), against the observed associations with AMD ($y$-axis). Model 1 including LDL diameter (LDL.D), and TG content in small HDL (S.HDL.TG) is shown in the main manuscript. The colour code shows: left) the $q$-statistic for outliers and right) Cook's distance for the influential points. Any genetic variant with $q$-value larger than 10 or Cook's distance larger than the median is marked by a label indicating the gene region.

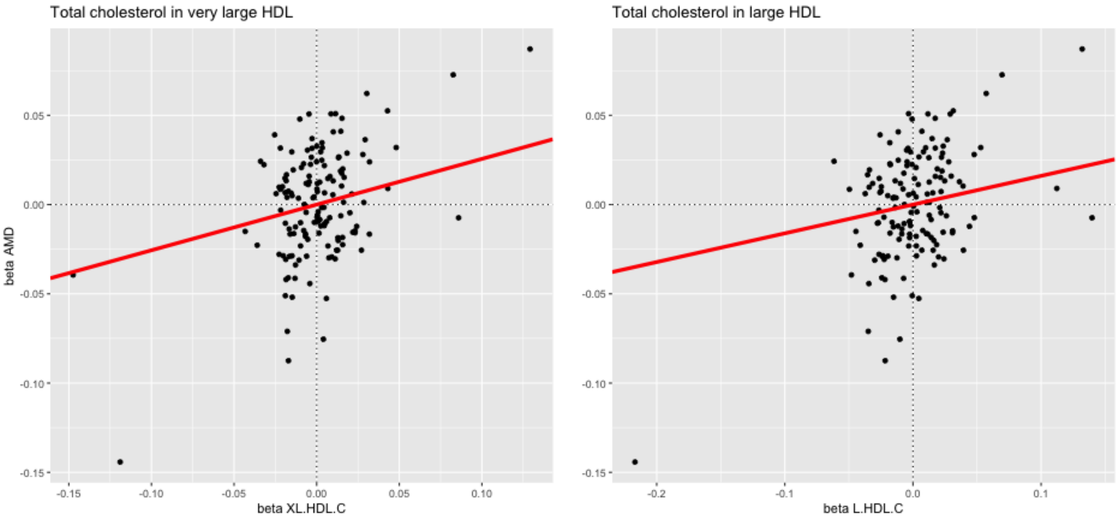

Figure 17: Scatterplot of association with A) XL.HDL.C B) L.HDL.C on the $x$-axis against the association with AMD $y$-axis after excluding the *LIPC*, *FUT2* and *APOE* gene regions. The model-averaged causal effect (MACE) of each risk factor on AMD is marked in red.

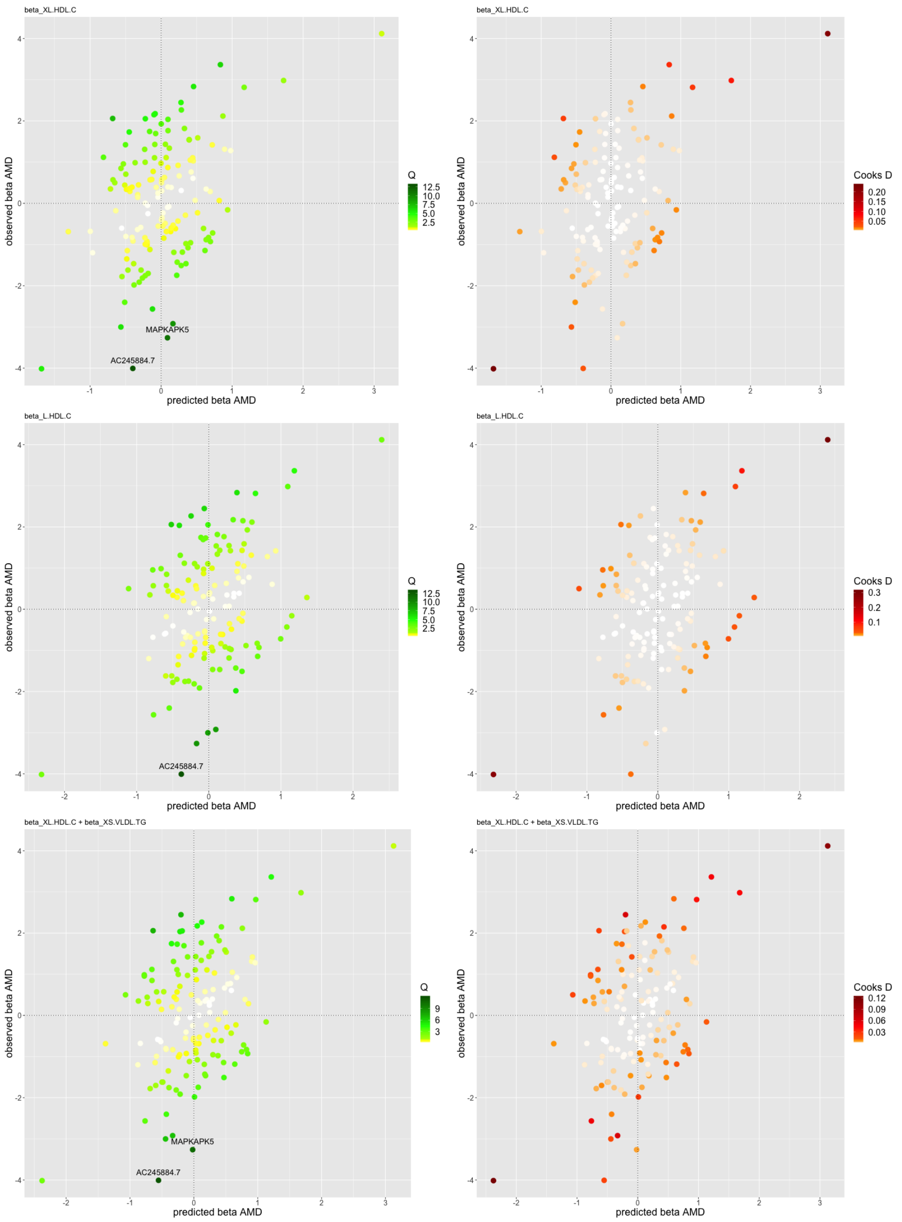

Figure 18: Diagnostic plots of the predicted associations with AMD ($x$-axis) based on the best individual model 1 (M1: XL.HDL.C), model 2 (M2: L.HDL.C), model 3 (M3: XL.HDL.C and XS.VLDL.TG against the observed associations with AMD ($y$-axis). The colour code shows: left) the $q$-statistic for outliers and right) Cook's distance for the influential points. Any genetic variant with $q$-value larger than 10 or Cook's distance larger than the median is marked by a label indicating the gene region. The *LIPC, FUT2* and *APOE* gene regions have been removed prior to this analysis.

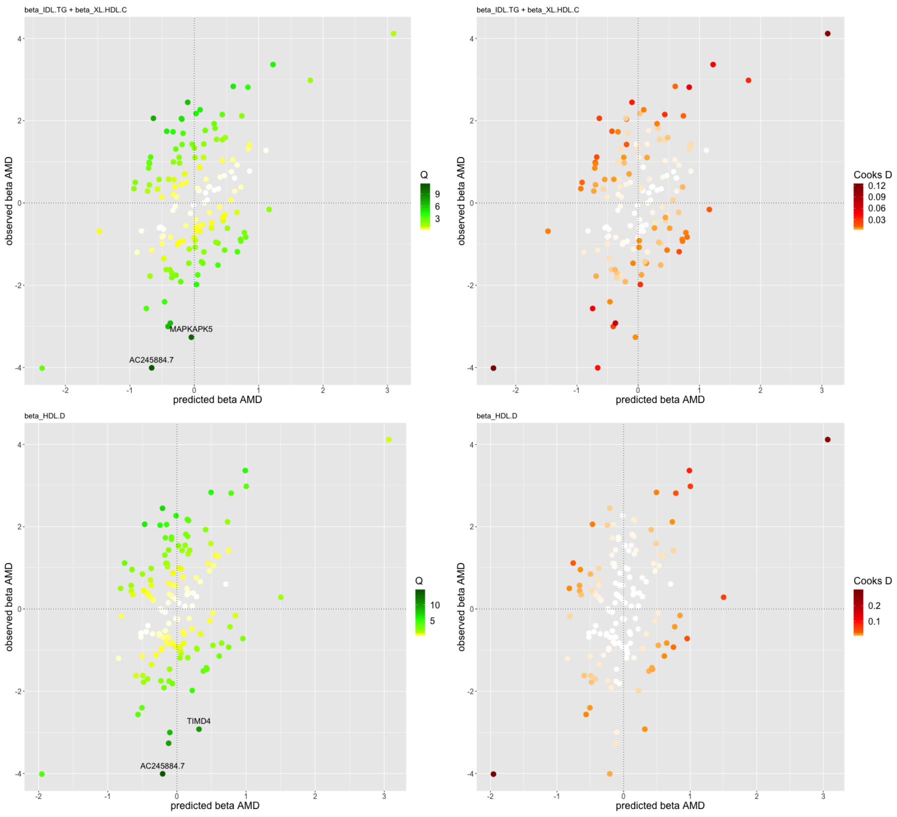

Figure 19: Diagnostic plots of the predicted associations with AMD ($x$-axis) based on the best individual model 4 (M4: IDL.TG and XL.HDL.C), model 5 (M5: HDL.D) against the observed associations with AMD ($y$-axis). The colour code shows: left) the $q$-statistic for outliers and right) Cook's distance for the influential points. Any genetic variant with $q$-value larger than 10 or Cook's distance larger than the median is marked by a label indicating the gene region. The *LIPC*, *FUT2* and *APOE* gene regions have been removed prior to this analysis.
