## Supplementary Note S1 for "Selecting causal risk factors from high-throughput experiments using multivariable Mendelian randomization"

### 1 Supplementary Note S1: Derivation of Bayes factors for a set of risk factors

In this note, we derive a closed form expression for the Bayes factor that quantifies the evidence for a particular model (one risk factor or set ofmultiple risk factors) to have a causal effect on the outcome compared to the Null model, which includes no risk factor and no intercept.

Building on the 2-sample MR approach [1] our work is based on summarised data, where genetic variants are used as instrumental variables. In univariable MR, we observe for each genetic variant $i = 1, ..., n$ the association of variant $i$ with the risk factor $X$ measured by the beta-coefficient $\beta_{X_i}$ from a univariable regression where the genetic variant $i$ is regressed on the risk factor $X$, and the association of variant $i$ with the outcome $Y$ measured by the beta-coefficient $\beta_{Y_i}$ where the genetic variant $i$ is regressed on the outcome $Y$, respectively. The causel effect $\theta$ of risk factor $X$ on $Y$ can be estimated using the IVW estimate or equivalently the following weighted regression without an intercept

$$\beta_{Y_i} = \theta \beta_{X_i} + \epsilon_i, \quad \epsilon_i \sim \mathcal{N}(0, \mathrm{se}(\beta_{Y_i})^2). \tag{1}$$

The same causal effect $\theta$ can be derived using a 2-stage least squares approach [2]. In fact, the beta-coefficients are estimates of the genetic association, but we omit the "hat" notation and treat the beta-coefficient as observations. A further assumption for this approach is that the genetic variants are independent (or uncorrelated) which can be controlled in the selection process of the genetic variants. Extension for correlated variants are for example described in [2].

In order to consider multiple risk factors jointly in one model multivariable MR was introduced in [3]. In the following, we consider $j = 1, ..., d$ risk factors. Assume $\boldsymbol{\beta}_\mathbf{X} = \{\beta_{X_1}, ..., \beta_{X_d}\}$ to be a matrix of dimension $n \times d$, where $d$ is the number of risk factors and $n$ is the number of genetic variants. Again each individual element $\beta_{X_{i,j}}$ of the predictor matrix is derived from a univariable regression where the genetic variant $i$ is regressed on the risk factor $X_j$. Multivariable MR can be cast as a weighted linear multivariable regression model

$$\beta_{Y_i} = \theta_1 \beta_{X_{i1}} + \theta_2 \beta_{X_{i2}} + \ldots + \theta_d \beta_{X_{id}} + \epsilon_i, \quad \epsilon_i \sim \mathcal{N}(0, \mathrm{se}(\beta_{Y_i})^2), \tag{2}$$

where the dependent variable is the association with the outcome $\beta_Y$ measured on $i = 1, ...., n$ instrumental variables and the predictors are the $j = 1, ..., d$ genetic associations with the $d$ risk factors. In matrix notation this can be written as

$$\beta_Y = \boldsymbol{\beta}_\mathbf{X} \boldsymbol{\theta} + \epsilon, \qquad \epsilon_i \sim \mathcal{N}(0, \mathrm{se}(\beta_{Y_i})^2). \tag{3}$$

In other words, the risk factors represent the variable space and the genetic variants used as instrumental variables are treated as observations. In practise, we standardise each observation of both, $\beta_{Y_i}$ and $\beta_{X_i}$ by dividing by $se(\beta_{Y_i})$ before the analysis and we assume in the following derivations that $\beta_Y$ and $\boldsymbol{\beta}_\mathbf{X}$ are standardised.

We use Bayes factors [4] in order to quantify the evidence for a particular model. By model we refer to the set of risk factors which have a causal effect on the outcome of interest. In order to formalise which risk factors are part of a specific model $M_{\boldsymbol{\gamma}}$ and consequently have a causal effect on the outcome we introduce a binary indicator $\boldsymbol{\gamma}$ of length $d$ that indicates which risk factors are selected and which ones are not

$$\gamma_j = \begin{cases} 1, & \text{if the } j\text{th risk factor is selected,} \\ 0 & \text{otherwise.} \end{cases} \tag{4}$$

The indicator $\boldsymbol{\gamma}$ encodes a specific regression model $M_{\boldsymbol{\gamma}}$ that includes the risk factors as indicated in $\boldsymbol{\gamma}$. Accordingly, we define $\boldsymbol{\beta}_{\mathbf{X}_{\boldsymbol{\gamma}}}$ as the design matrix of the risk factors included and $\boldsymbol{\theta}_{\boldsymbol{\gamma}}$ as the respective causal effects.

The computation of the Bayes factor for model $M_{\boldsymbol{\gamma}}$ against the Null model $M_0$, i.e. including no risk factor and no intercept, as presented in the Methods section of the main article requires two ingredients: First the marginal probability of $\beta_Y$ given $\boldsymbol{\beta}_{\mathbf{X}_{\boldsymbol{\gamma}}}$ of model $M_{\boldsymbol{\gamma}}$ and second, the marginal probability of $\beta_Y$ given the Null model $M_0$, which we derive as follows:

1. $p_{\boldsymbol{\gamma}}(\beta_Y \mid \boldsymbol{\beta}_{\mathbf{X}_{\boldsymbol{\gamma}}})$: the marginal probability of $\beta_Y$ given $\boldsymbol{\beta}_{\mathbf{X}_{\boldsymbol{\gamma}}}$

   In order to model the correlation between risk factors we base our likelihood on a multivariate Gaussian distribution

$$\beta_Y \mid \boldsymbol{\beta}_{\mathbf{X}_{\boldsymbol{\gamma}}}, \boldsymbol{\theta}_{\boldsymbol{\gamma}}, \tau \sim N(\boldsymbol{\beta}_{\mathbf{X}_{\boldsymbol{\gamma}}} \boldsymbol{\theta}_{\boldsymbol{\gamma}}, \frac{1}{\tau}). \tag{5}$$

   Following Servin and Stephens' $D_2$ prior [5] we use the following conjugate prior assumptions for the causal effects $\boldsymbol{\theta}_{\boldsymbol{\gamma}}$, the residual $\epsilon$ and the precision $\tau$

$$\begin{aligned} \boldsymbol{\theta}_{\boldsymbol{\gamma}} \mid \tau &\sim N(0, \boldsymbol{\nu}/\tau), \\ \epsilon &\sim N(0, \frac{1}{\tau}), \\ \tau &\sim \Gamma(\kappa/2, \lambda/2), \end{aligned} \tag{6}$$

   where $A \mid B$ is defined as $A$ conditional on $B$. Further we can derive analytically the joint posterior distribution for $\boldsymbol{\theta}_{\boldsymbol{\gamma}}$ and $\tau$ as

$$\begin{aligned} \tau \mid \beta_Y, \boldsymbol{\beta}_{\mathbf{X}_{\boldsymbol{\gamma}}} &\sim \Gamma((n+\kappa)/2, 1/2(\beta_Y^t \beta_Y - \boldsymbol{\Theta}^{\mathbf{t}} \boldsymbol{\Omega}^{-\mathbf{1}} \boldsymbol{\Theta} + \lambda)), \\ \boldsymbol{\theta}_{\boldsymbol{\gamma}} \mid \beta_Y, \boldsymbol{\beta}_{\mathbf{X}_{\boldsymbol{\gamma}}}, \tau &\sim N(\boldsymbol{\Theta}, \frac{1}{\tau} \boldsymbol{\Omega}), \end{aligned}$$

   where

$$\underbrace{\boldsymbol{\Theta}}_{d \times 1} = \underbrace{\boldsymbol{\Omega}}_{d \times d} \underbrace{\boldsymbol{\beta}_{\mathbf{X}_{\boldsymbol{\gamma}}}{}^t}_{d \times n} \underbrace{\beta_Y}_{n \times 1}, \tag{7}$$

$$\boldsymbol{\Omega} = \underbrace{(\boldsymbol{\nu}^{-1} + \boldsymbol{\beta}^{\mathbf{t}}_{\mathbf{X}_{\boldsymbol{\gamma}}} \boldsymbol{\beta}_{\mathbf{X}_{\boldsymbol{\gamma}}})^{-1}}_{d \times d}. \tag{8}$$

Next we integrate out the causal effects $\boldsymbol{\theta_\gamma}$. To begin with, we sort the equation so that the integral contains only terms dependent on $\boldsymbol{\theta_\gamma}$

$$
\begin{aligned}
p_{\boldsymbol{\gamma}}(\beta_Y \mid \ \boldsymbol{\beta}_{\mathbf{X}_\gamma}, \tau) &= \int_{-\infty}^{\infty} \frac{p_{\boldsymbol{\gamma}}(\beta_Y \mid \boldsymbol{\beta}_{\mathbf{X}_\gamma}, \boldsymbol{\theta_\gamma}, \tau) p_{\boldsymbol{\gamma}}(\boldsymbol{\theta_\gamma} \mid \tau)}{p_{\boldsymbol{\gamma}}(\boldsymbol{\theta_\gamma} \mid \beta_Y, \boldsymbol{\beta}_{\mathbf{X}_\gamma}, \tau)} \delta\boldsymbol{\theta_\gamma} \\
&= \int_{-\infty}^{\infty} \frac{(2\pi)^{-\frac{n}{2}} \tau^{\frac{n}{2}} \exp\left\{-\frac{\tau}{2}(\beta_Y - \boldsymbol{\beta}_{\mathbf{X}_\gamma}\boldsymbol{\theta_\gamma})^t(\beta_Y - \boldsymbol{\beta}_{\mathbf{X}_\gamma}\boldsymbol{\theta_\gamma})\right\}}{(2\pi)^{-\frac{1}{2}} \frac{\|\boldsymbol{\Omega}\|^{-1/2}}{\|\tau\|^{-1/2}} \exp\left\{-\frac{\tau}{2}(\boldsymbol{\theta_\gamma} - \boldsymbol{\Theta})^t \boldsymbol{\Omega}^{-1}(\boldsymbol{\theta_\gamma} - \boldsymbol{\Theta})\right\}} \\
&\quad \times (2\pi)^{-\frac{1}{2}} \frac{\|\boldsymbol{\nu}\|^{-1/2}}{\|\tau\|^{-1/2}} \exp\left\{-\frac{\tau}{2\boldsymbol{\nu}} \boldsymbol{\theta}_{\boldsymbol{\gamma}}^{\mathbf{t}} \boldsymbol{\theta_\gamma}\right\} \delta\boldsymbol{\theta_\gamma} \\
&= (2\pi)^{-\frac{n}{2}} \tau^{\frac{n}{2}} \frac{\|\boldsymbol{\Omega}\|^{1/2}}{\|\nu\|^{1/2}} \exp\left\{-\frac{\tau}{2}(\beta_Y^t \beta_Y - \boldsymbol{\Theta}^{\mathbf{t}} \boldsymbol{\Omega}^{\mathbf{-1}} \boldsymbol{\Theta})\right\} \\
&\quad \times \int_{-\infty}^{\infty} \exp\left\{-\frac{\tau}{2}\left(-2\boldsymbol{\theta}_{\boldsymbol{\gamma}}^{\mathbf{t}} \boldsymbol{\beta}_{\mathbf{X}_\gamma}^{\mathbf{t}} \beta_Y + \boldsymbol{\theta}_{\boldsymbol{\gamma}}^{\mathbf{t}} (\boldsymbol{\beta}_{\mathbf{X}_\gamma}^{\mathbf{t}} \boldsymbol{\beta}_{\mathbf{X}_\gamma} - \frac{1}{\boldsymbol{\nu}}) \boldsymbol{\theta}_\gamma - \right.\right. \\
&\qquad\qquad \left.\left. \boldsymbol{\theta}_{\boldsymbol{\gamma}}^{\mathbf{t}} \boldsymbol{\Omega}^{\mathbf{-1}} \boldsymbol{\theta}_\gamma + 2\boldsymbol{\theta}_{\boldsymbol{\gamma}}^{\mathbf{t}} \boldsymbol{\Omega}^{\mathbf{-1}} \boldsymbol{\Theta}\right)\right\} \delta\boldsymbol{\theta}_\gamma,
\end{aligned}
$$

where $\|\mathbf{X}\|$ denotes the determinant of a matrix $\mathbf{X}$ and $\infty$ infinity. Note that the final line in the integral can be simplified to

$$
-2\boldsymbol{\theta}_{\boldsymbol{\gamma}}^{\mathbf{t}}(\mathbf{A} - \mathbf{D}) + \boldsymbol{\theta}_{\boldsymbol{\gamma}}^{\mathbf{t}}(\mathbf{B} - \mathbf{C})\boldsymbol{\theta}_\gamma, \tag{9}
$$

where

$$
\begin{aligned}
\mathbf{A} &= \boldsymbol{\beta}_{\mathbf{X}_\gamma}^{\mathbf{t}} \beta_Y \\
\mathbf{B} &= (\boldsymbol{\beta}_{\mathbf{X}_\gamma}^{\mathbf{t}} \boldsymbol{\beta}_{\mathbf{X}_\gamma} - \frac{1}{\boldsymbol{\nu}}) \\
\mathbf{C} &= \boldsymbol{\Omega}^{\mathbf{-1}} \\
\mathbf{D} &= \boldsymbol{\Omega}^{\mathbf{-1}} \boldsymbol{\Theta}
\end{aligned} \tag{10}
$$

By completing the square in $\boldsymbol{\theta_\gamma}$ and integrating out $\boldsymbol{\theta_\gamma}$ the final integral equals 1.

Overall, this simplifies to

$$
p_{\boldsymbol{\gamma}}(\beta_Y \mid \ \boldsymbol{\beta}_{\mathbf{X}_\gamma}, \tau) \quad = (2\pi)^{-\frac{n}{2}} \tau^{\frac{n}{2}} \frac{\|\boldsymbol{\Omega}\|^{1/2}}{\|\boldsymbol{\nu}\|^{1/2}} \exp\left\{-\frac{\tau}{2}(\beta_Y^t \beta_Y - \boldsymbol{\Theta}^{\mathbf{t}} \boldsymbol{\Omega}^{\mathbf{-1}} \boldsymbol{\Theta})\right\} \tag{11}
$$

Next we integrate out the precision $\tau$

$$
\begin{aligned}
p_{\boldsymbol{\gamma}}(\beta_Y \mid \ \boldsymbol{\beta}_{\mathbf{X}_\gamma}) &= \int_0^{\infty} p_{\boldsymbol{\gamma}}(\beta_Y \mid \ \boldsymbol{\beta}_{\mathbf{X}_\gamma}, \tau) p(\tau) \delta\tau \qquad (12) \\
&= (2\pi)^{-\frac{n}{2}} \frac{\|\boldsymbol{\Omega}\|^{1/2}}{\|\boldsymbol{\nu}\|^{1/2}} \\
&\quad \times \int_0^{\infty} \tau^{\frac{(n+\kappa)}{2}-1} \exp\left\{-\frac{1}{2}(\beta_Y^t \beta_Y - \boldsymbol{\Theta}^{\mathbf{t}} \boldsymbol{\Omega}^{\mathbf{-1}} \boldsymbol{\Theta} + \lambda)\tau\right\} \delta\tau.
\end{aligned}
$$

The above integral is the normalisation constant of a Gamma distribution with shape $\alpha = \frac{(n+\kappa)}{2}$ and rate $\beta = \frac{1}{2}(\beta_Y^t \beta_Y - \mathbf{\Theta^t \Omega^{-1} \Theta} + \lambda)$. Thus the above simplifies exactly to

$$p_{\boldsymbol{\gamma}}(\beta_Y \mid \ \boldsymbol{\beta}_{\mathbf{X}_{\boldsymbol{\gamma}}}) \quad = (2\pi)^{-\frac{n}{2}} \frac{\|\mathbf{\Omega}\|^{1/2}}{\|\boldsymbol{\nu}\|^{1/2}} (\tfrac{\lambda}{2})^{\frac{\kappa}{2}} \frac{\Gamma(\frac{n+\kappa}{2})}{\Gamma(\frac{\kappa}{2})} \tag{13}$$

$$\times \left\{\tfrac{1}{2}(\beta_Y^t \beta_Y - \mathbf{\Theta^t \Omega^{-1} \Theta} + \lambda)\right\}^{\frac{-(n+\kappa)}{2}}. \tag{14}$$

2. $p_0(\beta_Y)$: the marginal probability of $\beta_Y$ given the Null model $M_0$

Next, we derive the marginal probability of the Null model, i.e. where no risk factor and no intercept is included. Under the Null we assume

$$\beta_Y \mid \frac{1}{\tau} \sim N(0, \frac{1}{\tau}) \tag{15}$$

with an expectation fixed at zero, which is a consequence of the missing intercept.

First, we integrate out the precision $\tau$

$$\begin{aligned} p_0(\beta_Y) &= \int_0^\infty p_0(\beta_Y \mid \tau) p(\tau) \delta\tau \\ &= (2\pi)^{-\frac{n}{2}} \int_0^\infty \tau^{\frac{(n+\kappa)}{2}-1} \exp\left\{-\frac{1}{2}(\beta_Y^t \beta_Y + \lambda)\tau\right\} \delta\tau. \end{aligned} \tag{16}$$

Again the above integral is the normalisation constant of a Gamma distribution with shape $\alpha = \frac{(n+\kappa)}{2}$ and rate $\beta_0 = \frac{1}{2}(\beta_Y^t \beta_Y + \lambda)$. Thus the above simplifies to

$$p_0(\beta_Y) = (2\pi)^{-\frac{n}{2}} (\frac{\lambda}{2})^{\frac{\kappa}{2}} \frac{\Gamma(\frac{n+\kappa}{2})}{\Gamma(\frac{\kappa}{2})} \left(\frac{1}{2}(\beta_Y^t \beta_Y + \lambda)\right)^{-\frac{(n+\kappa)}{2}}. \tag{17}$$

The Bayes factor for model $M_{\boldsymbol{\gamma}}$ against $M_0$ is defined as the ratio of the marginal probability of $\beta_Y$ given model $M_{\boldsymbol{\gamma}}$ (13) over the marginal probability of $\beta_Y$ given the Null model (17)

$$\begin{aligned} BF(M_{\boldsymbol{\gamma}}) &= \frac{p_{\boldsymbol{\gamma}}(\beta_Y \mid \boldsymbol{\beta}_{\mathbf{X}_{\boldsymbol{\gamma}}})}{p_0(\beta_Y)} \\ &= \frac{\frac{\|\mathbf{\Omega}\|^{1/2}}{\|\nu\|^{1/2}} \left(\frac{1}{2}(\beta_Y^t \beta_Y - \mathbf{\Theta^t \Omega^{-1} \Theta} + \lambda)\right)^{-(n+\kappa)/2}}{\left(\frac{1}{2}(\beta_Y^t \beta_Y + \lambda)\right)^{-(n+\kappa)/2}} \\ &= \frac{\|\mathbf{\Omega}\|^{1/2}}{\|\boldsymbol{\nu}\|^{1/2}} \left(\frac{\beta_Y^t \beta_Y - \mathbf{\Theta^t \Omega^{-1} \Theta} + \lambda}{\beta_Y^t \beta_Y + \lambda}\right)^{-(n+\kappa)/2}. \end{aligned} \tag{18}$$

Next we consider the limit as $\kappa$, $\lambda \to 0$. $\kappa$ and $\lambda$ are the shape and scale parameter of the Gamma Distribution, which is the conjugate distribution for precision. In the limiting case the expectation of the error precision would converge towards a point mass at zero. A precision that converges to zero translates into an error variance that converges to infinity. Thus the limiting case represents a dominant error noise and no variance explained by the model, which is a conservative prior assumption. Moreover, the limit $\lambda \to 0$ leads to the invariance property of the posterior for $\theta$, ie the posterior of $\theta$ changes appropriately with shifts and scaling (for example inverse-variance weighting) operations on $\beta_y$.

In limit, the Bayes Factor simplifies to the following closed form expression

$$BF(M_\gamma) = \frac{\|\boldsymbol{\Omega}\|^{1/2}}{\|\nu\|^{1/2}} \left( \frac{\beta_Y^t \beta_Y - \boldsymbol{\Theta^t \Omega^{-1} \Theta}}{\beta_Y^t \beta_Y} \right)^{-n/2}. \tag{19}$$

These Bayes factors are then used in the model averaging to quantify the evidence for a model and together with the prior are used to evaluate which model or set of risk factors has the largest support to have a causal effect on the outcome.
